# Change in brain asymmetry reflects level of acute alcohol intoxication and impacts on inhibitory control

**DOI:** 10.1101/2023.01.10.523048

**Authors:** Julien Dubois, Ryan M. Field, Sami Jawhar, Austin Jewison, Erin M. Koch, Zahra M. Aghajan, Naomi Miller, Katherine L. Perdue, Moriah Taylor

## Abstract

Alcohol is one of the most commonly used substances and frequently abused, yet little is known about the neural underpinnings driving variability in inhibitory control performance after ingesting alcohol. This study was a single-blind, placebo-controlled, randomized design with participants (N=48) completing three study visits. At each visit participants received one of three alcohol doses; namely, a placebo dose (equivalent Blood Alcohol Concentration (BAC) = 0.00%), a low dose of alcohol (target BAC=0.04%), or a moderate dose of alcohol (target BAC=0.08%). To measure inhibitory control, participants completed a Go/No-go task paradigm twice during each study visit, once immediately before dosing and once after, while their brain activity was measured with time-domain functional near-infrared spectroscopy (TD-fNIRS). BAC and subjective effects of alcohol were also assessed. We report decreased behavioral performance for the moderate dose of alcohol, but not the low or placebo doses. We observed right lateralized inhibitory prefrontal activity during go-no-go blocks, consistent with prior literature. Using standard and novel metrics of lateralization, we were able to significantly differentiate between all doses. Lastly, we demonstrate that these metrics are not only related to behavioral performance during inhibitory control, but also provide complementary information to the legal gold standard of intoxication (i.e. BAC).

## Introduction

Acute alcohol intoxication is widely known to impact cognitive functions including executive control and psychomotor functioning^1,2^. Some cognitive functions are highly sensitive to alcohol and are impaired at relatively low doses, while other functions appear to be preserved even at higher doses of alcohol^3,4^. Inhibitory control, the ability to stop a prepotent motor response after it has been initiated, appears to be particularly sensitive to the influence of alcohol^5^. Impairment of the inhibitory control system has been implicated in behaviors associated with alcohol intoxication such as, reckless driving, impulsivity, increased aggression, and an inclination towards risky actions^6–8^.

In a laboratory setting, inhibitory control is often measured with a Go/No-go or Stop Signal task^9^. In a standard Go/No-go paradigm participants are asked to respond rapidly to the timed presentation of “Go stimuli” and withhold their motor response to less frequent “No-go stimuli”. Most behavioral studies using the Go/No-go inhibitory control paradigms have shown that after drinking alcohol, participants had worse performance and altered reaction times^1,10,11^.

While the behavioral effects of alcohol consumption are relatively consistent, the impact of alcohol on brain activity during inhibitory control tasks is less clear. Functional magnetic resonance imaging (fMRI) studies have shown that the prefrontal regions are implicated in inhibitory control. In particular, the right prefrontal cortex plays a critical role in this function, revealed as a distinct asymmetry in brain activity between right and left prefrontal regions when individuals are engaged in no-go conditions of the task^12–14^. Considering the acute effect of alcohol on the prefrontal cortex, the fMRI literature has been conflicting^15^. For example, some studies have shown mixed effects^16^ while others have shown decreases^11,17,18^ in prefrontal activity during inhibitory control tasks. This inconsistency could be a result of varied dosing and subsequent varied intoxication levels across studies, yet few studies have implemented more than one dose of alcohol^18,19^. Beyond prefrontal activation, to our knowledge, only one fMRI study and one fNIRS study have directly looked at prefrontal asymmetry (lateralization) under acute alcohol intoxication and the relationship to disinhibition^11,20^.

Functional near-infrared spectroscopy (fNIRS), which also measures brain hemodynamic activity, is a rapidly growing brain imaging technique. While fNIRS imaging is limited to cortical regions, it has advantages over fMRI including: ease-of-use, portability/wearability, safety, and being more cost-effective. In this vein, fNIRS provides a unique opportunity to examine the brain under the influence of alcohol^21^ given its ecological validity, safety and reduced susceptibility to movement artifacts. Indeed, prior fNIRS literature has replicated the findings of prefrontal involvement in inhibitory control^22–24^. However, only a handful of fNIRS studies have explored the effect of acute alcohol intoxication on the brain^20,25^. For example, it has been reported that prefrontal lateralization changes acutely after alcohol consumption and the same group provided preliminary evidence that the change in prefrontal lateralization during a Go/No-go task is correlated with task performance^20^.

Some studies have looked at dose-dependent effects of alcohol on the brain^18,19^, and others have focused on prefrontal activation and lateralization during inhibitory control tasks following acute intoxication^11,20^. However, further research is required to examine whether the prefrontal dynamics exhibit a dose-dependent pattern and how these dynamics relate to behavior. In this study, we go beyond prior studies and fully explore the interrelations between prefrontal activation patterns and changes in inhibitory behavior at varied alcohol intoxication levels. Functional brain activity was measured using a time-domain functional near-infrared spectroscopy (TD-NIRS) system, Kernel Flow1^26^, while participants performed a version of the Go/No-go task before and after three separate single-blind alcohol dosing sessions (placebo, low, and moderate).

Here, we report group-level dose-dependent changes in behavior, subjective feelings of intoxication, and brain activity. In doing so, we quantify observed changes in prefrontal asymmetry by not only considering a standard measure of brain lateralization, but also introduce novel measures of lateralization that are time-locked and more granular. Furthermore, we explore the relationship between task performance, alcohol intoxication levels (measured breath alcohol content (BrAC) converted to equivalent blood alcohol content (BAC)), and neural factors (prefrontal lateralization), while also considering demographic information. This analysis elucidates the role that lateralized prefrontal activity plays in inhibitory control under the influence of acute alcohol intoxication.

## Results

### Experimental Design and Data Collection

Participants were 48 healthy individuals (23 male, age (mean ± s.d.)=32.63 ± 9.90 years, age range=38 years; Methods; Supplementary Table 1) who completed multiple study visits during which they performed a series of inhibitory control tasks before and after they consumed a beverage containing either a placebo, low, or moderate dose of alcohol (Fig. 1A, Methods). The inhibitory control task consisted of blocks with low demand of control (go-only blocks) and high demand of control (go-no-go blocks)(Fig. 1b, Methods).

**Figure 1:**
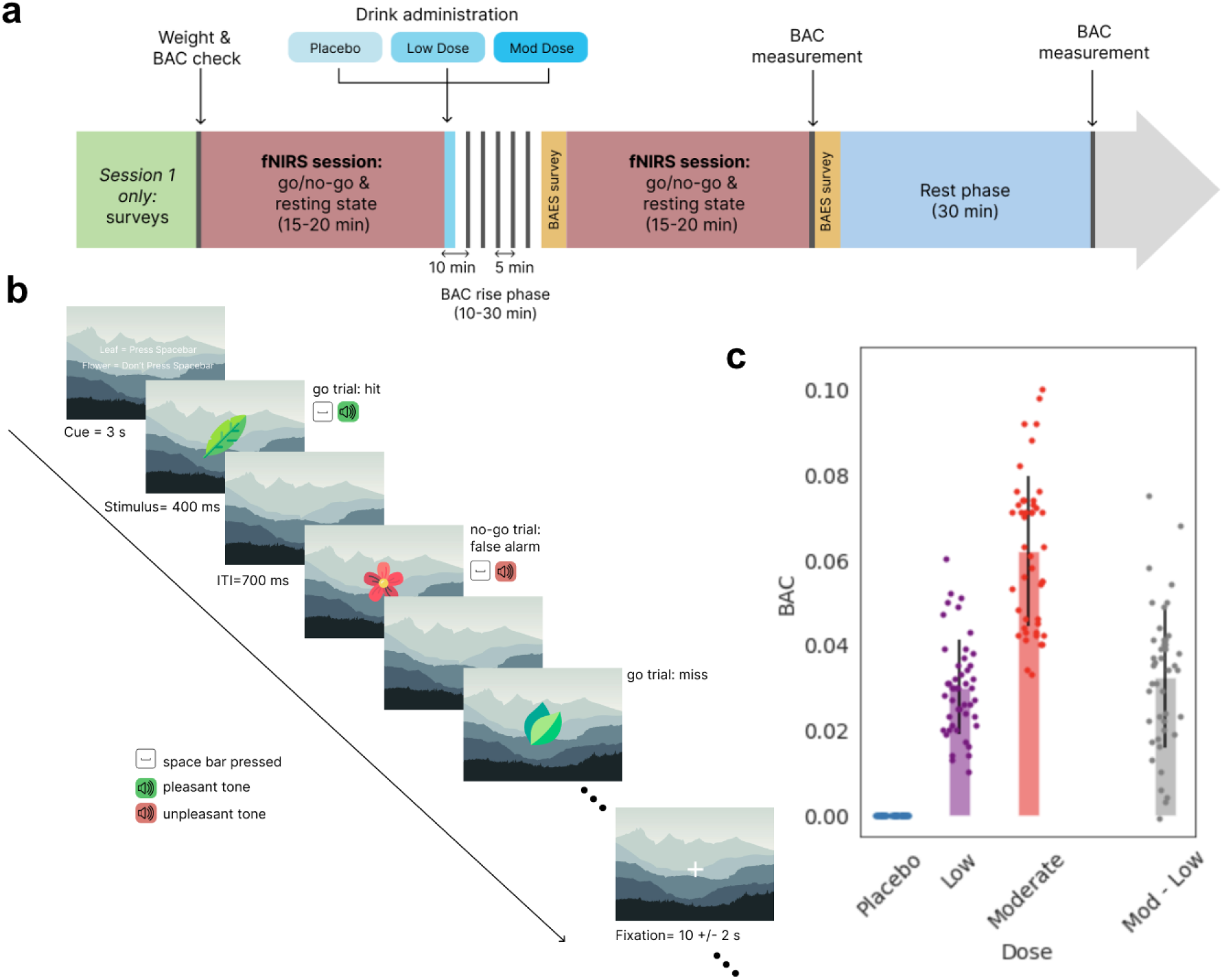
Overview of study design. **a.** The structure of each experimental session: participants performed an inhibitory control task (Go/No-go) while wearing the Kernel Flow1 headset. Next, depending on the predetermined randomized order, they were given a beverage with placebo (zero), low, or moderate doses of alcohol. Following alcohol consumption, there was a wait period during which BAC was measured every five minutes using a breathalyzer until the target BAC level was reached (10-30 minutes; gray vertical lines indicate BAC measurements). After the wait period, participants performed another round of the Go/No-go task. **b.** Schematic of the Go/No-go block structure. A few representative trials at the start of a go-no-go block are shown. **c.** BAC values for each participant (each dot) measured using a breathalyzer after the BAC rise phase and before the second Go/No-go task. BAC values after the moderate dose were significantly higher compared to those after the low dose of alcohol (paired t-test p=2.15×10^−16^; shown are mean±s.d.).

We targeted a BAC of 0.04 and 0.08 for low and moderate sessions respectively. When BACs were measured using a breathalyzer (Methods), we found that on average participants had lower than intended targeted values. However, when given the moderate dose of alcohol, participants consistently had a higher BAC compared to the lower dose of alcohol (Fig. 1C). During the tasks, hemodynamics signals were captured using the Kernel Flow1 headset capable of recording whole-head time-domain functional near-infrared spectroscopy (TD-NIRS) data^26^.

To capture the subjective feeling of intoxication, we asked participants to complete the Biphasic Alcohol Effects Scale^27^ before and after their post-dose task as well as predict the number of standard drinks they thought they were given (Methods). In agreement with the literature, subjective feelings of intoxication were different between dosing sessions. A repeated measures ANOVA indicated a significant effect of dose on the sedative score, stimulant score, and drink prediction (p<0.05 for all). For all subjective measures, there was a significant difference between all pairs of doses (paired t-test p<0.05 for all; Supplementary Fig. 1). In summary, subjects showed differences in their measured level of intoxication (BAC) and subjective feelings of intoxication across dosing sessions, laying the proper foundation for the behavioral and neural analyses described below.

### Effect of alcohol dose on inhibitory control behavioral metrics

In order to assess the effect of alcohol dose on inhibitory control, we first computed participants’ behavioral performance on the Go/No-go tasks before and after alcohol consumption (henceforth referred to as pre and post respectively). These metrics include the subject’s performance (measured as d-prime in go-no-go blocks) and their reaction times (during go-no-go blocks).

We noted that pre performance and post performance were strongly correlated for all doses, but this relationship deteriorated progressively with higher doses (placebo: r=0.75; low: r=0.51; moderate: r=0.48)(Fig. 2a). In fact, as intoxication level increased from placebo to low and then low to moderate, there was a systematic decrease in the slope of the line-of-best fit as well as a monotonic reduction in the correlation and significance of the relationship. We observed that for the moderate dose most of the points fall below the line of unity indicating a drop in d-prime between pre and post runs.

**Figure 2:**
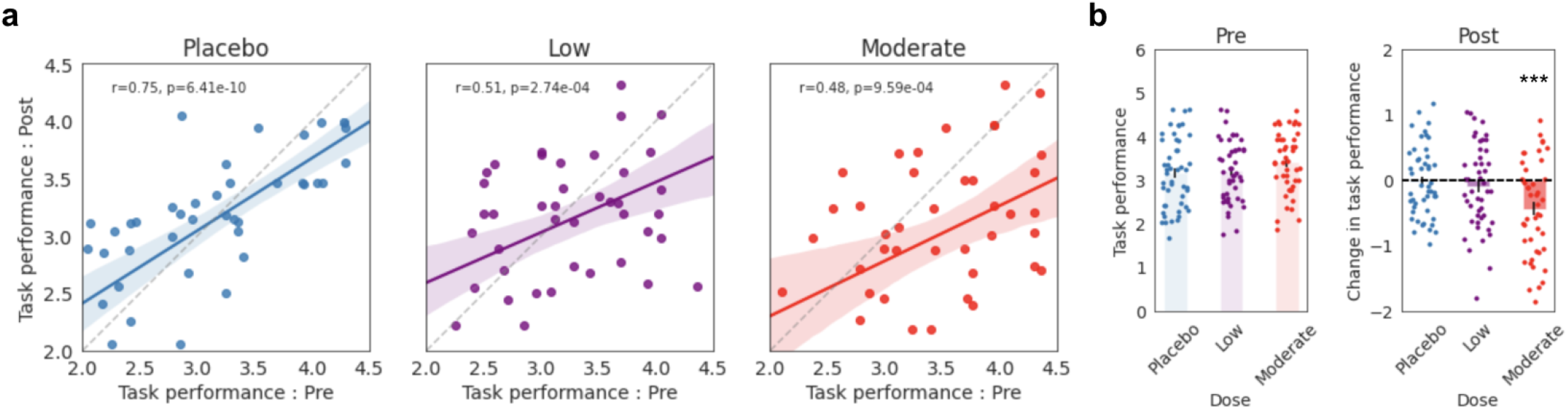
Main effect of alcohol level on task performance as measured by d-prime. **a.** Pre Go/No-go performance is plotted against post Go/No-go performance for placebo (left, blue), low (middle, purple), and moderate (right, red) levels of intoxication. Solid lines depict the line-of-best-fit and the shaded regions show the 95% confidence intervals. Dashed lines indicate the line of unity. Task performance is computed as the d-prime in the go-no-go blocks. **b.** Left) Colored bars show the average d-prime (task performance) across the pre tasks for placebo, low, and moderate sessions separately. Right) The change in performance, measured as post d-prime - pre d-prime are shown for the different doses. Colored dots show individual subject measures (bar plots: mean±s.e.m). Behavioral performance was significantly impaired in the post task compared to the pre task for the moderate dose (paired t-test p=2.35×10^−4^).

To quantify these observations, we ran a two-way repeated measures ANOVA to model each behavioral metric using factors pre/post, alcohol dose, and their interaction term. We found that the interaction between pre/post and dose was a significant factor in d-prime (p=0.003), but not the reaction time (p=0.200). For reaction time, only the main effect of pre/post was deemed significant (p=0.037; Supplementary Fig. 2). There was no relationship with dose, which is inconsistent with previous findings indicating slower reaction times with intoxication^18,28^. Additionally, we found that in the moderate dose, there was a significant reduction in post performance (d-prime) compared to pre (paired t-test p=2.35×10^−4^)(Fig. 2b). When only considering the post performance, we found a significant difference between the placebo and moderate doses (paired t-test p<0.05). On the other hand, we found no significant behavioral change in low dose sessions. These results suggest that behavioral deficits, while more prominent at higher doses, are not reliable indicators of intoxication, especially at low doses of alcohol. Thus, we wondered whether brain activity could offer more robust and granular measures of intoxication.

### Alcohol dose and different patterns of prefrontal brain activation

While participants were performing both pre- and post-dosing Go/No-go tasks, hemodynamic brain activity was acquired using Kernel Flow1^26^. Raw data, i.e. the photons’ distribution of time of flights (DToFs) for two wavelengths (690 and 850nm) underwent standard preprocessing steps that include obtaining DToF moments (sum, mean, and variance) and converting their variations into relative concentrations of oxygenated hemoglobin (HbO) and deoxygenated hemoglobin (HbR), as described previously^26,29^ (Methods).

To investigate brain activation patterns associated with our Go/No-go inhibition task, we employed a GLM framework, and modeled the activity of each channel as a function of the two block types (go-only and go-no-go) as well as other commonly used task-irrelevant regressors (Methods). We performed this analysis for each chromophore independently. Full-head brain activity patterns were identified for each dosing session (Supplementary Fig. 3). While these results implicate a number of brain regions in the Go/No-go task (including prefrontal regions and anterior temporal regions), we limited our analyses by only using the channels formed between sources and detectors in the prefrontal region (HbO: Fig. 3a-b, HbR: Fig. 3c-d; Methods). This was inspired by prior literature that points towards this region’s role in inhibitory control^14,20^. During the pre-dose task, we found increased HbO during go-no-go blocks (compared to go-only blocks) in the right prefrontal cortex and reduced HbO in the left anterior prefrontal cortex. The measured difference in HbO between left and right prefrontal cortex, termed lateralization, is in agreement with prior literature^12,20^ (Fig. 3a).

**Figure 3:**
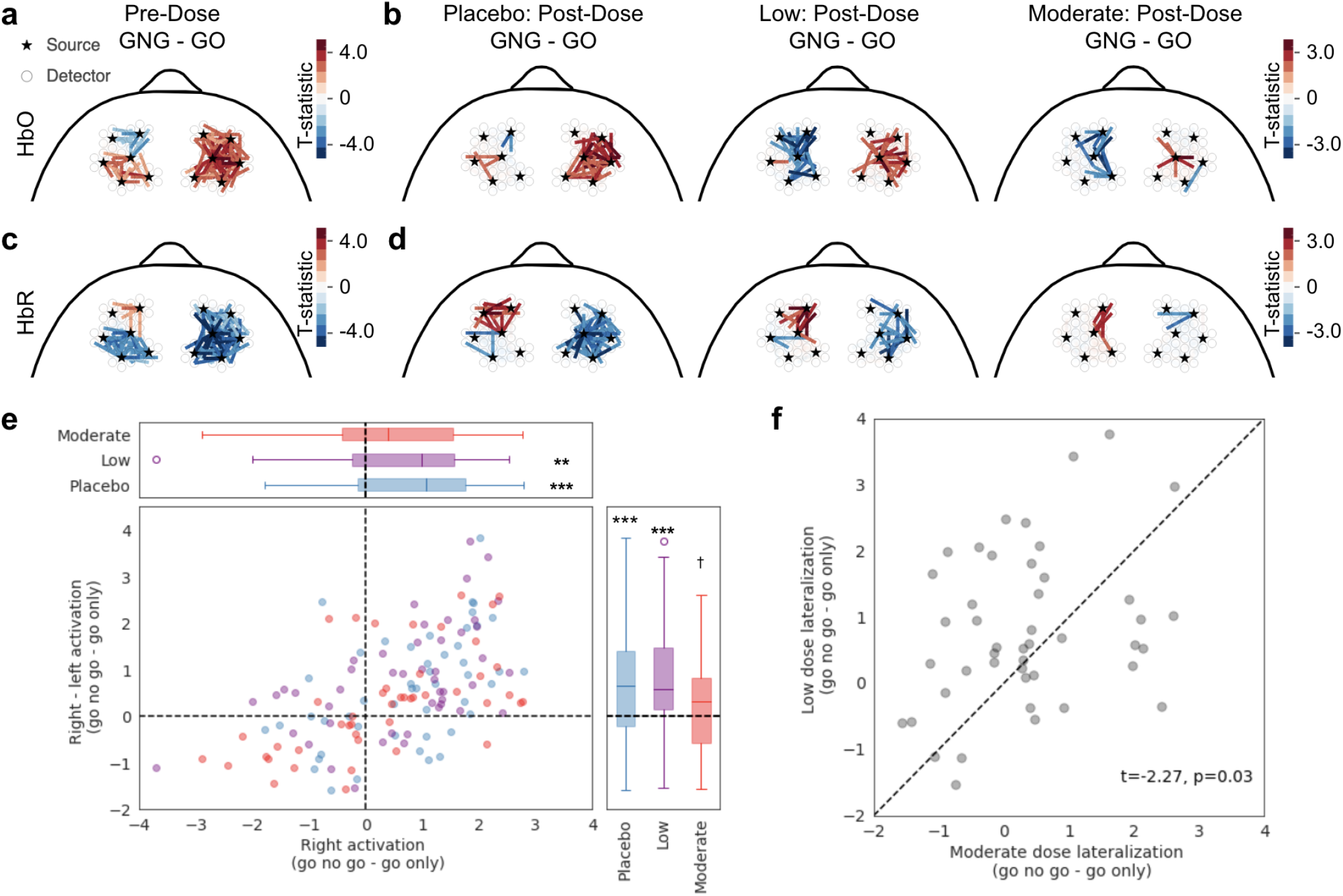
Prefrontal brain activation during an inhibitory control task under the influence of different alcohol doses as quantified by generalized linear models. **a.** HbO in pre-dose Go/No-go tasks showed increased right prefrontal activation during go-no-go (GNG) blocks compared to the go-only (GO) blocks (as indicated by warmer colors on the right side). This activation is accompanied by anterior deactivation in the left prefrontal cortex (indicated by cooler colors). Each line is an individual channel and only significant channels at p<0.05 (uncorrected) are displayed. **b.** Same as (a), but showing activation patterns for placebo (left), low (middle), and moderate (right) doses of alcohol during the post Go/No-go task. Different dosing sessions exhibited different strengths and patterns of lateralization. **c–d.** Same as (a–b) except for HbR. In all subpanels, the color scale indicates the t-statistic. **e.** Average right - average left brain activity (GLM effect size for go-no-go - go-only contrast) plotted against average right brain activity. Dose-separated distributions are displayed via box plots along the top and right margins. The distributions were compared against zero using one-sample t-test (** and *** indicate significance at p<0.01 and p<0.001 respectively; † indicate trends towards significance). **f.** Right - left average GLM effect size (GLM-based lateralization) was significantly higher for low-versus moderate-dose (shown are paired t-test statistics). Dashed line indicates the line of unity.

Looking across post-dose GLM results, it appeared that the strength and spread of the right prefrontal activity during go-no-go blocks (compared to go-only), as well as the overall patterns of lateralization varied according to alcohol dose (Figure 3b). When we repeated the GLM analysis for HbR, we observed generally complementary (to HbO) patterns of task-related activity(Fig 3c, d). In order to quantify the observed right prefrontal activity during go-no-go blocks, contrasted with go-only, for each participant we averaged the effect size (coefficients from the GLM analysis) of all the channels in the right prefrontal region. First, the population distribution of average right prefrontal activity was significantly positive for both placebo and low doses (placebo: t=4.57, p=3.52×10^−5^; low: t=3.18, p=0.002; one-sample t-test), but this increase was absent in the moderate dose (Figure 3e top). Second, to consider GLM-based lateralization, we repeated this analysis on the difference between average right and left prefrontal activation. Here a positive difference indicates right dominance (in go-no-go compared to go-only condition). Placebo and low doses had significant right lateralization (placebo: t=3.64, p=6.74×10^−4^; low: t=4.75, p=2.02×10^−5^; one-sample t-test); while the moderate dose showed a weaker trend (t=1.88, p =0.067; one-sample t-test) (Figure 3e right). It is interesting to point out that lateralization was strongest for the low dose, suggesting the potential presence of compensatory mechanisms. Last, we performed a direct comparison of the GLM-based lateralization across the two alcohol doses and found that lateralization in the moderate dose was significantly lower (on average) compared to the low dose (moderate vs low: t=-2.27, p=0.028; paired t-test), indicating that with this analysis we can differentiate between the different doses of alcohol (Fig. 3f).

Because the aforementioned analysis contrasted averaged responses between the go-no-go and go-only blocks, it might obscure changes in the time courses of activation patterns within each condition. Thus, to obtain a more granular view into the temporal dynamics of lateralization, we sought to define lateralization metrics that took into account the time courses of activity averaged over occurrences of each block type, within each hemisphere (Fig. 4).

**Figure 4:**
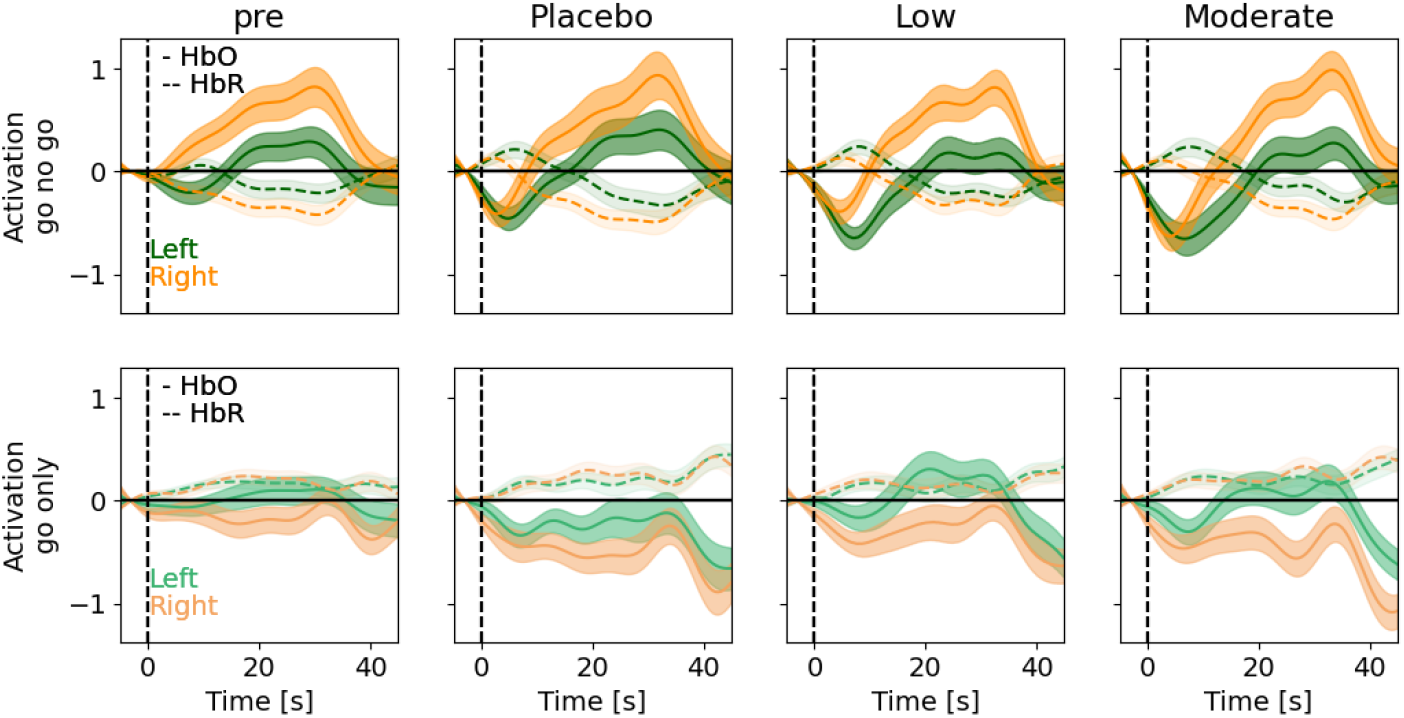
Time courses of prefrontal lateralization for different alcohol doses. Epoched time course of left (green) and right (orange) prefrontal activity for go-no-go blocks (top) and go-only blocks (bottom) separated by session. From left to right, pre-dose and placebo-, low-, and moderate-dose time courses are shown (mean±s.e.m). Solid and dashed lines represent HbO and HbR respectively. Note the (expected) opposite trends in the two chromophores.

The epoched HbO/HbR time courses during go-no-go blocks (Fig. 4 top) and go-only blocks (Fig. 4 bottom) exhibited the expected opposite trends for the two chromophores. We noted that during the go-no-go blocks, the right prefrontal HbO was consistently higher than the left prefrontal HbO, while the opposite was true for the go-only blocks. To quantify this, we computed another measure of lateralization (termed direction metric) as the averaged signed difference between right and left activity, such that a positive value indicates right-lateralization (Methods). This reversal of lateralization between the two block types, as seen in HbO (go-no-go: t=8.91, p=6.50×10^−17^; go-only: t=-5.84, p=1.43×10^−8^) was notably missing from HbR (go-no-go: t=-7.43, p=1.30×10^−12^; go-only: t=0.86, p=0.39). A likely explanation is that the concentration of oxygenated hemoglobin results in larger signals and patterns of activity tasks as opposed to concentrations of deoxygenated hemoglobin^30^.

It has been suggested that total hemoglobin (HbT = HbO + HbR) is more robust in mapping cerebral activation (i.e. less contaminated by superficial signals)^30^. Indeed, HbT exhibited the most distinct time courses across doses, while also demonstrating the aforementioned lateralization (Fig. 5a). Consequently, HbT will be used as a proxy for brain activity in the remaining analyses. At the population level, the direction metric was significantly positive (i.e. right lateralization) for the go-no-go blocks (t=5.14, p=5.08×10^−7^) and significantly negative (i.e. left lateralization) for the go-only blocks (t=-5.36, p=1.73×10^−7^)(Fig. 5b). In addition to differentiating between task conditions (go-only and go-no-go blocks), when comparing the direction metric computed during go-no-go blocks across post-dose conditions, we found a statistically discernible difference between the placebo/low doses (p = 0.037) (Fig. 5c). Of interest, the DToF mean moment of 850nm alone struck a clear resemblance to the HbT time courses (Supplementary Fig. 4a). This may not be surprising given that the mean moment captures changes in deeper tissue^31^, and while being more sensitive to changes in oxy-hemoglobin, it also incorporates information about deoxy-hemoglobin^31–33^. The mean moment (of 850 nm) also revealed complementary results when considering the direction metric (Supplementary Fig. 4b,d).

**Figure 5:**
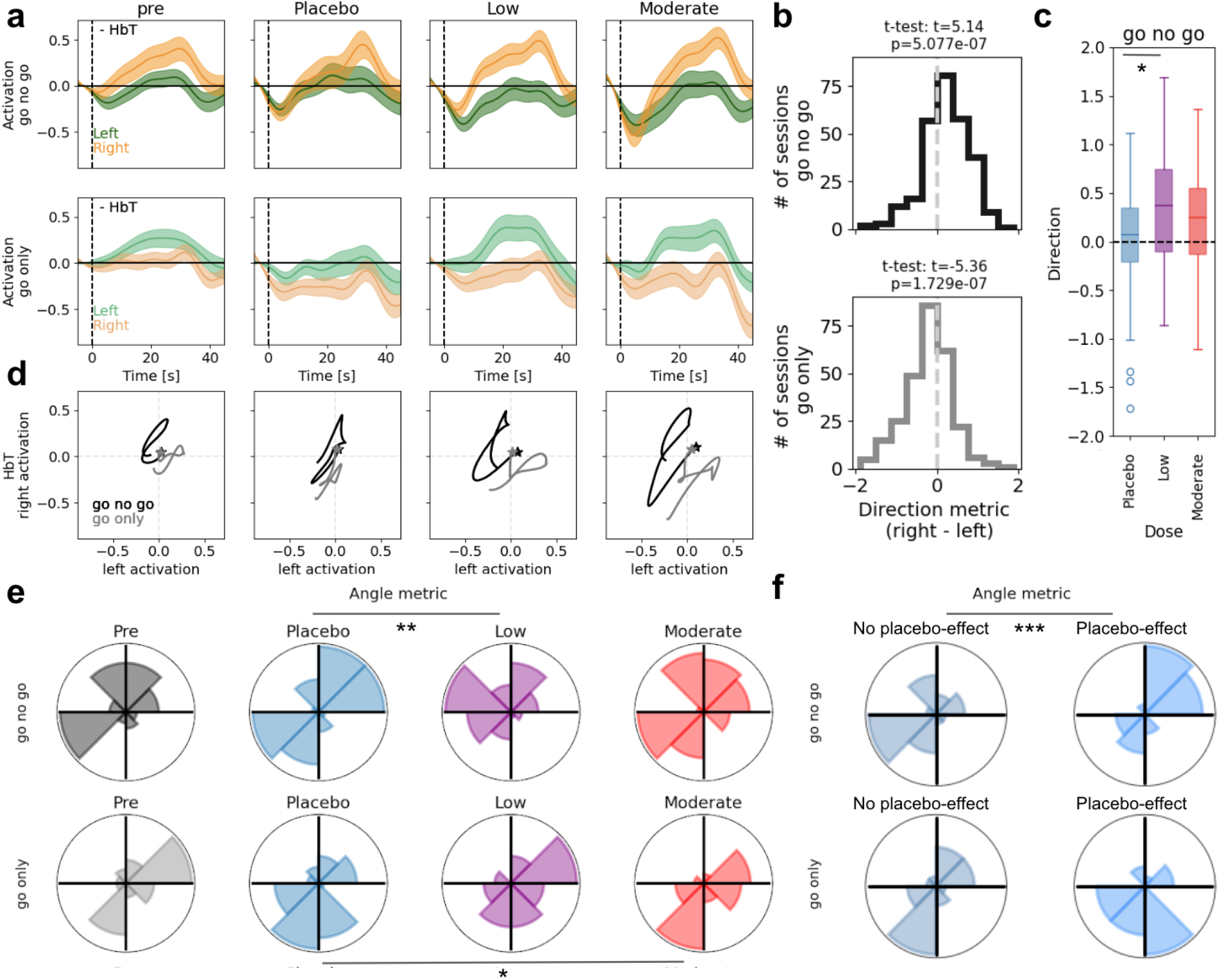
Altered patterns of prefrontal lateralization with different alcohol doses. **a.** Block-averaged time courses of left (green) and right (orange) prefrontal HbT for go-no-go blocks (top) and go-only blocks (bottom) separated by session. From left to right, pre-dose and placebo-, low-, and moderate-dose time courses are shown (mean±s.e.m). **b.** Population distribution of lateralization (direction metric) for go-no-go (top; black) and go-only (bottom; gray) blocks across all participants and sessions. Note the right lateralization for go-no-go blocks (shift towards positive values, t-test: p=5.08×10^−7^) and left lateralization for go-only blocks (shift towards negative values, t-test: p=1.73×10^−7^). **c.** The distributions of the direction metric for go-no-go blocks across post-dose sessions (* significant difference between placebo and low dose: paired t-test p=0.037). **d.** Population average parametric curves showing the trajectory of right prefrontal vs. left prefrontal activity through time for go-only (gray) and go-no-go (black) blocks, split by pre-dose and session type, as in (a). Stars indicate initial time points. **e.** The distributions of the angle metric, split by pre-dose and session type, as in (a). * and ** indicate significant deviation from uniformity (at *p* < 0.05 and p < 0.01 respectively; rayleigh test) when computing pairwise differences in the angle metric between dosing sessions. **f.** Distributions of the angle metric, split by no placebo-effect/placebo-effect (left/right) and go-no-go blocks/ go-only blocks (top/bottom). *** indicates a significant difference in the population mean in the go-no-go condition (p=9.80×10^−5^; Watson-Williams test) comparing no placebo-effect (n=23, circular mean=176.50 deg) and placebo effect (n=23, circular mean=34.45 deg).

The direction metric, while informative, is still a coarse measure of lateralization that does not take into account time-locked covariability between left and right prefrontal activity. To further characterize how the time courses of left and right prefrontal activity co-varied, we inspected the parametric curves defined by the right versus left prefrontal HbT through time within each block type separately (Fig. 5d). We noted that the trajectories were qualitatively different not only between the block types, but also across different dosing sessions. To quantify these differences, we computed the average trajectory, which is defined by an angle and magnitude in the two dimensional parametric space (x- and y-axes corresponding to the left and right prefrontal activity respectively, Methods). Importantly, these novel metrics offered additional information about absolute activation (or deactivation), beyond the direction metric.

The distributions of angle metric as seen in Fig. 5e, highlighted a difference between the placebo and the two alcohol doses. To measure this, we computed the pairwise difference in the angle metric (between doses) and compared the resulting distribution (angle-difference distribution) against a uniform distribution. We found that in the go-no-go condition, the angle-difference distribution between placebo and low doses was significantly discernable from a uniform distribution (rayleigh test: p=0.002, mean difference = −46.52 deg). In the go-only condition, the significant contrast was found between placebo and moderate doses (rayleigh test: p=0.019, mean difference = −27.13 deg). Pairwise statistical comparison of the magnitude metric revealed a trend towards significance between placebo/moderate doses (paired t-test: p=0.061) in the go-only blocks, with low alcohol dose showing a larger magnitude metric than placebo (Supplementary Fig. 5a).

Could these lateralization metrics differentiate participants who felt a placebo-effect from those that did not? In our population, almost half of the participants (n = 23) reported a placebo effect (Methods). We wondered if the placebo-effect was driving the bimodal nature of the angle metric distribution in the placebo dose (Fig. 5e). Indeed, we found that the angle metric exhibited significantly different distributions between those who had a placebo-effect and those who did not (Watson Williams test: p=9.80×10^−5^)(Fig. 5f). Although, we cannot fully rule out the effect of the session order in which a placebo drink was administered as there was a strong overlap with those who received placebo in their first visit and those who reported having received alcohol.

These lateralization metrics, taken together, provide a foundation for distinguishing intoxicated brain activity. Interestingly, when using only the 850nm mean moment we were able to replicate these findings, and we have the addition of statistically significant differences between the low and moderate doses (Supplementary Fig. 4c-e, Supplementary Fig5b). In the remaining analyses we will investigate to what degree the change in behavior can be captured by lateralization metrics while considering confounds, such as demographics, BAC, and subjective reports.

### The importance of prefrontal brain dynamics in determining inhibitory control

As shown earlier, post performance was strongly correlated with pre performance, but the strength of this relationship decreased with increasing levels of intoxication (Fig. 2a). It is possible that other variables may explain the change in performance from pre to post. Variables of potential interest include: direct measure of intoxication, or the BAC values; and brain measures, i.e. the different lateralization metrics. As expected, behavior was more impaired with higher levels of BAC (Fig. 6a top; pearson r=-0.29, p=5.24×10^−4^). In agreement with prior literature^20^, we found a positive correlation between the direction metric and change in performance (Fig. 6a bottom; pearson r=0.22, p=7.47×10^−3^) indicating that stronger right lateralization resulted in better inhibitory control. A combination of the angle and magnitude metrics, i.e. transformation from polar to cartesian coordinates, also showed a significant correlation with the change in behavior when considering the sine component (magnitude x sin(angle)) (Fig. 6a middle; pearson r=0.19, p=2.63×10^−2^), but not the cosine component. It is important to note that we did not find a relationship between BAC and any of the above lateralization metrics (p>0.1 for both direction and angle metrics), underscoring that not only do these metrics contain information about behavior, but that this information is complementary (to BAC) and not overlapping.

**Figure 6:**
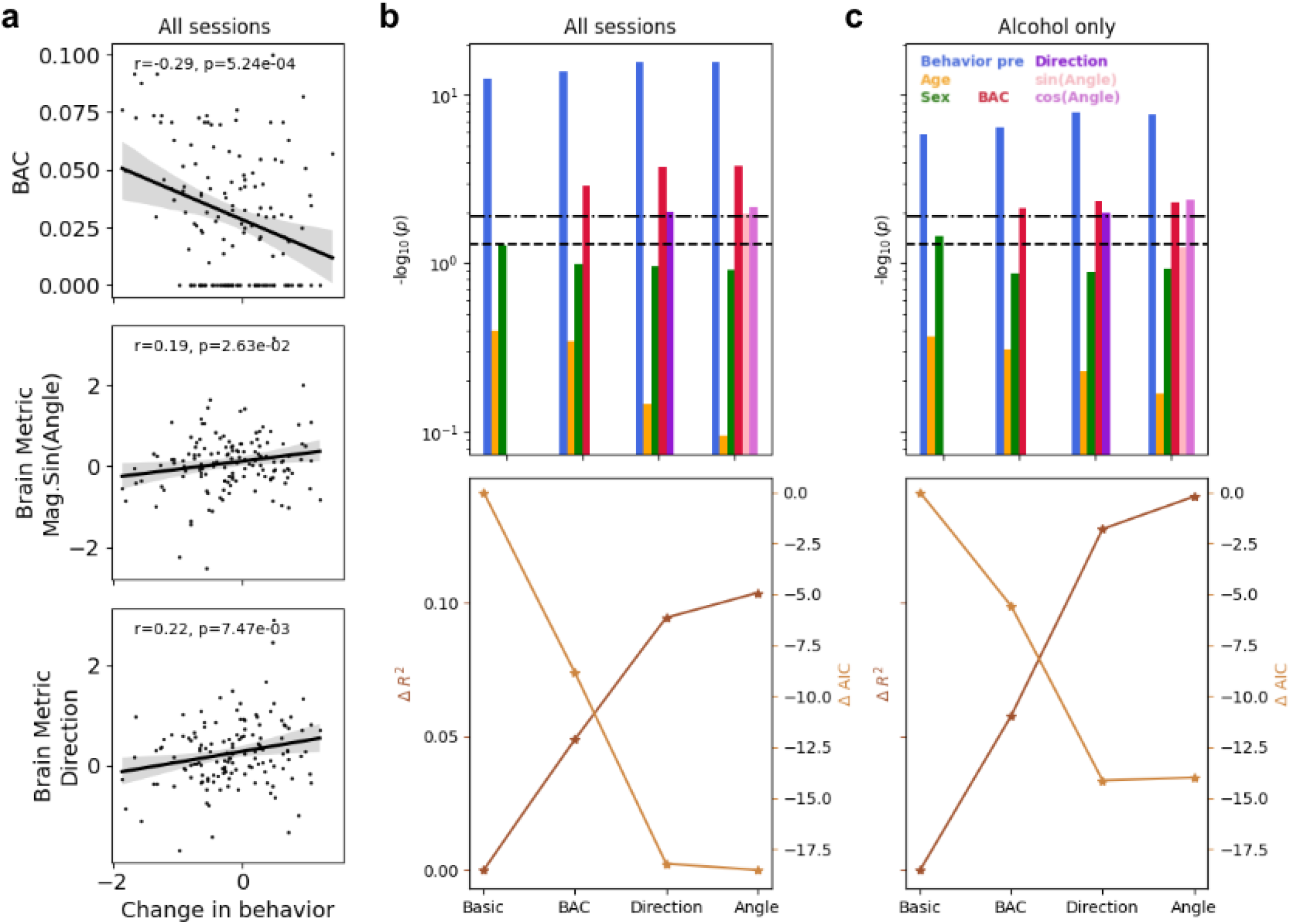
Lateralization metrics improved the prediction of behavioral performance in addition to measured BAC. **a.** The change in behavioral performance as measured by post d-prime - pre d-prime was significantly correlated with BAC (top; pearson r=-0.29, p=5.24×10^−4^), magnitude x sin(angle) (middle; pearson r=0.19, p=2.63×10^−2^), and direction metric (bottom; pearson r=0.22, p=7.47×10^−3^). **b.** Top) Statistical significance of the different factors in the four OLS models (four blocks shown on the x-axis) in predicting the post behavioral performance are reported as −log_10_ (p-value). We build off of the three factors in the basic model (pre behavioral performance: blue, age: yellow, sex: green), and include additional variables in separate OLS models: 1) BAC model included the direct measure of intoxication (BAC values, red) in addition to the basic model; 2) Direction model consisted of the direction metric (dark purple) in addition to the BAC model variables; 3) Angle model included the weighted sine and cosine of the angle metric (pink and light purple respectively) in addition to the BAC model variables. Note that both BAC values and lateralization metrics were significant factors in predicting post performance. Dashed lines indicate a significance level of 0.05 and 0.05/4 (after correcting for multiple comparisons using Bonferroni correction; Methods). Bottom) Overall OLS model performances for the four models shown in terms of the change in R-squared values and the change in AIC (lower AIC values indicate better performance). **c.** Same as (b) but when only considering the sessions with alcohol consumption (i.e. low and moderate doses).

To tease apart the contribution of different parameters to post behavior, we used an Ordinary Least Squares (OLS) approach (Methods) and compared model performances when considering different sets of input parameters. We started with a baseline model, where post performance was predicted as a linear combination of pre performance (Fig. 2a), sex, and age. As expected, pre performance was a significant factor in predicting post performance, but both age and sex factors failed to pass significance thresholds (Fig. 6b top). Next, we included the direct measure of intoxication, i.e. BAC values, as an additional term in the model and found BAC to be a significant predictor of performance in conjunction with pre behavior (Fig. 6b top; termed BAC model). The model performance showed an overall improvement (Fig. 6b bottom) both in terms of increased R-squared and decreased Akaike information criterion (AIC), which is important to consider to avoid model overfitting with the addition of new variables (Methods).

Next, we wondered whether the addition of the lateralization metrics, namely the direction and angle metrics, would improve our model performance in explaining behavior. As such, two additional models were considered in which the direction metric and angle metric were added to the BAC model independently (Methods). Note that in order to take into account the circular nature of the angle metric in our linear model, we included both the sine and cosine terms (weighted by the magnitude of the vector). Both direction and angle metrics proved to be statistically relevant to the model (Fig. 6b top) and, in fact, both of these models outperformed the BAC model in terms of model fit and explained variance (Fig. 6b bottom). Curiously, when subjective measures of intoxication were added to the BAC model, they did not reach statistical significance and, in fact, decreased the model performance.

We repeated the above analysis while considering only sessions that involved alcohol consumption (i.e. low and moderate) doses. The motivation to perform this analysis was to ensure that placebo sessions were not driving the model performance and diluting the importance of alcohol-related features. Namely, when considering all dosing sessions, including placebo, the distribution of BACs contains an over representation of 0.00 (one-third of the sessions). Therefore, considering only dosing sessions provides a mathematically sound and fairer comparison of the effect of BAC and lateralization metrics on inhibitory control. When comparing the performance of the different OLS models on alcohol-only sessions, we observed that the lateralization metrics (i.e. the direction and angle metrics) performed comparable to BAC in terms of feature importance. The improvement in model performance by the addition of the lateralization metrics was even more pronounced when considering only alcohol sessions (Fig 6b). Highlighting this, the addition of the angle metric increased explained variance by almost 15% beyond the basic model, while also claiming a lower AIC than the BAC model.

Taken together, these analyses suggest that the brain lateralization metrics provide complementary information to the legal gold standard of intoxication (i.e. BAC), and may offer more robust predictive power, when considering behavioral performance during inhibitory control. Following suit, the brain-behavior relationship as well as the above-mentioned descriptive model–when redone with 850nm mean moment–matched these results (Supplementary Fig. 6).

## Discussion

Inhibitory control is a complex cognitive function, involving many coordinated brain regions, and neurobiology that is not yet fully understood. Likewise, the mechanisms underlying disruptions of the inhibitory control system, either due to a neurobiological disorder, such as ADHD^34^, or ingestion of a substance, such as alcohol, remain unclear. Nevertheless, a large body of work implicates the right prefrontal cortex as having a causal role in both inhibitory control and disinhibition^12–14,35,36^. Within the brain imaging community, fMRI studies have reported changes in behavior and prefrontal activity during performance of an inhibitory task (Go/No-go or Stop-Signal) when participants are under the influence of acute alcohol intoxication^11,17,18^. Recently, fNIRS has emerged as a motion-tolerant, portable, and cheaper alternative to fMRI. In fact, fNIRs seems particularly well-suited for studies involving intoxicated subjects; however, only a few have used this technology in this domain^20,25^. In this study, we used TD-fNIRS to measure brain activity during a version of the Go/No-go task, implementing a multi-dose single-blind procedure, targeting low intoxication (BAC=0.04) and moderate intoxication (BAC=0.08). This study design allowed us to build on prior literature and investigate changes in inhibitory behavior, neural activity, and their inter-relations across multiple doses of alcohol, further elucidating the role of prefrontal dynamics.

We found that inhibitory behavior, as measured by the Go/No-go task, was only affected at moderate doses of alcohol, such that performance was significantly worse after alcohol consumption compared to before. Correspondingly, we found that post-moderate performance was significantly worse than post-placebo performance, but not post-low performance. These results indicate that moderate levels of acute alcohol intoxication significantly change subjects’ ability to discriminate and respond accurately to go and no-go stimuli. In this version of the task, we found that at low levels of intoxication subjects performed comparably to pre-dose and post-placebo sessions. Furthermore, we only observed a significant decrease in reaction times in the placebo sessions (post-ingestion compared to pre-ingestion) suggesting that the unimpaired individual can execute the task faster without compromising performance. While we did not replicate prior findings of change in reaction times across different intoxication levels^1,18,28^, we hypothesize that this may be due to the difficulty level of our specific implementation of the task. A larger implication of these results is that behavioral deficits, while prominent at higher levels of intoxication, are not always reliable indicators of intoxication at lower doses.

We examined whether brain activity during the inhibitory control task could provide more reliable measures of intoxication. As a first step, we confirmed the importance of prefrontal regions during the Go/No-go task^12–14^, as observed in both our GLM and epoched analyses when performed prior to alcohol consumption. Moreover, we not only found a right hemispheric dominance during the go-no-go blocks—consistent with prior literature^12–14^—but also found a reversal of lateralization during the go-only blocks. This finding is expected given the known predominance of control network regions on the right and default mode network regions on the left side of the prefrontal cortex^37^, as the cognitive demand of the go-only condition is minimal.

Because we observed a reliable change in performance at moderate levels of intoxication, we expected to see complementary disruptive changes in prefrontal brain activation. Indeed, it was only in the moderate dose that the expected activations patterns (both in terms of right activity and lateralization) disappeared in the GLM results. Furthermore, the moderate dose–when compared directly to the low dose of alcohol–showed significantly less GLM-based lateralization. At low levels of intoxication, task performance was unaffected, which could manifest as: 1. Unaltered activity (intoxication not strong enough), or 2. Changes in brain activity (possibly compensatory) that precede behavioral deficits. Qualitatively, we observed different patterns of significant prefrontal brain activity between low- and placebo-doses, yet, both showed GLM-based right lateralization–pointing towards a potential compensatory mechanism. To further assess differences between placebo and alcohol doses, we investigated metrics computed over the epoched time courses. While the direction metric distinguished between placebo and low-dose, our novel time-locked angle metric provided statistical discernibility between the placebo dose and both alcohol dosing sessions. To our knowledge, this is the most comprehensive comparison of dose-dependent changes in brain activity to date.

It is well-known that alcohol impairs inhibitory control, but the underlying contributing factors, neural and otherwise, remain unclear. In fact, our society uses BAC as the legal standard for impairment, even though subjective feelings of intoxication and behavioral performance across individuals can vary greatly when BAC is similar, a phenomenon often referred to as tolerance. In our data, pairwise correlations revealed a significant relationship between the change in behavioral performance and BAC (direct measure of intoxication), as well as both the direction and angle lateralization metrics (brain measures of intoxication). The correlation between the direction metric and performance is in agreement with Tsujii et. al^20^, although only one alcohol dose was considered in that study. We further gauged the relative importance of these factors, while controlling for other subject-level variables (sex, age), by employing successive OLS models and analyzing the statistical significance of each feature as well as the overall model performance. Our analyses showed that brain measures of lateralization were significant predictors of behavior and, when taken into account, they led to the highest performing models. One plausible explanation is that impairment state, mentioned above as tolerance, may be reflected in brain activity providing a more robust measure of intoxication than BAC values. It is important to note that we found no relationship between BAC and prefrontal lateralization, which supports the hypothesis that prefrontal lateralization can measure inhibitory impairment beyond physical intoxication.

It is worth noting that the current study has multiple limitations. First, there was an overlap in BAC values across dosing sessions. As a result, our group-level statistics included individuals with similar levels of BAC across low and moderate dosing sessions. In addition, half of our participants experienced a placebo-effect with corresponding changes in brain activity. Together, these confounds likely had a negative impact on our repeated measure statistics, some of which would not survive corrections for multiple comparisons. Second, it is difficult to administer a believable placebo dose of alcohol^15^. However, we followed protocols in the literature^38,39^ to circumvent these difficulties and improve the single-blind nature of the task design with half of the participants believing they received an alcohol dose. Third, for digestibility and practical reasons, we did not consider the full-breadth of subject-level factors in our modeling analysis. One that deserves mentioning is the handedness of the participants, as there have been reports of atypical lateralization in the left-handed population^40^. We did not include handedness in our model because we only had 5 left-handed participants. While we expect no direct effect of handedness on behavior, we hypothesize that including handedness could indirectly improve model performance by accounting for a potential reversal of prefrontal asymmetry; this remains to be explored in future studies. Last, it is likely that alcohol consumption leads to changes in blood flow^15^, which could be a potential confound in our analyses. While we did not observe a relationship between BAC and brain metrics, we nevertheless controlled for this by including measured BAC in our models. Further, we would not expect blood flow changes to be different between the two hemispheres and, therefore, this confound can be remedied by the fact that our neural analyses primarily involve metrics computing cross-hemispheric activation. Additionally, these metrics do not rely on strict adherence to canonical hemodynamic response functions and therefore may flexibly capture the full extent of changes in the hemodynamic response^15^.

There are multiple advantages of using a TD-fNIRS system for such investigations. Importantly, TD-fNIRS enhances depth sensitivity beyond the more commonly used continuous-wave (CW)-fNIRS, as it captures the arrival times of photons. Late arriving photons are more likely to have reached the brain than photons with earlier arrival times, which likely have only interacted with superficial tissues. The change in light absorption in the tissue at different depths can thereby be captured in the moments of DToFs (e.g. mean, variance). In fact, others have demonstrated the feasibility of using moments from a single wavelength in a TD-fNIRS system to detect brain activity^31^. With this in mind, in addition to performing analyses on chromophore concentrations, we explored the mean moment of the 850nm wavelength. With this single wavelength, we were able to identify dose-dependent differences in brain activity, and reproduce the relationship between brain metrics and behavior. This study, therefore, corroborates the proposition that a single wavelength from a TD-fNIRS system can provide measures of cerebral activation (under normal and altered states). This is critical for adapting fNIRS to applications that require faster implementation speed and lower cost, such as real-time neural feedback for brain-computer interfaces.

In short, this study provides critical evidence that dose-dependent changes in prefrontal brain activity can be measured with TD-fNIRS. We introduced novel metrics for measuring brain asymmetry that retain information about time-sensitive cross-hemispheric variability and absolute changes in brain activity. A highlight of this study is that we fully explored the extent to which changes in prefrontal lateralization can explain changes in task performance, by considering other potentially impactful features. The culmination of our study highlights the powerfulness and robustness of brain activity in explaining behaviorally relevant measures of intoxicated individuals. This underscores the importance of recording neural signals not just in academic laboratories seeking to understand neural mechanisms, but also in clinics, and pharmacological research.

## Methods

### Participants and screening procedures

Forty eight healthy participants (23 male, age=32.63 ± 9.90, age range=38) completed repeated study visits in a within-subject study design within three weeks (mean=5.67 days between visits, median=3.5, s.d.=6.16). Inclusion criteria were adults between the ages of 21–65, self-reported drinking at least twice a month, and body weight between 100 and 250 pounds. Exclusion criteria were current self-reported substance abuse disorder (excluding caffeine, tobacco, and cannabis) and a score of 2 or higher on the CAGE questionnaire^41^, medium risk or higher score on the Alcohol Use Disorders Identification Test (AUDIT)^42^, current or past enrollment in a substance use treatment program, current use of medications that are contraindicated with alcohol, known allergies to alcohol, pregnant or nursing, history of or current serious mental illness, neurological disorders, liver or cardiovascular diseases. Data was collected from an additional two participants but excluded from analysis for inability to perform the Go/No-go task (n=1) and failure to respond to alcohol dosing (e.g., BAC remained as 0 after low dose, n=1). Of the 48 included participants, 46 completed three study visits and 2 completed two study visits.

Participants signed the informed consent electronically in accordance with the IRB protocol (Advarra #Pro00063785). Participants received monetary compensation for their time, effort, and travel expenses. Participants were instructed and reminded not to drink alcohol within 24 hours of each study visit or eat 3 hours prior to each study visit. Participants were also required to arrange transportation after each study visit for safety.

### Study design

The study was a single-blind, placebo-controlled, randomized design with participants completing three study visits within three weeks (Fig. 1). Participants received either a placebo, low dose of alcohol (target BAC=0.04%), or moderate dose of alcohol (target BAC=0.08%) with the placebo visit always occuring in the first or second visit, and visit orders counterbalanced and randomly assigned over participants. To measure impulsivity and inhibitory control, participants completed a Go/No-go task paradigm twice during each study visit, once immediately before dosing and once after, while their brain activity was measured with Kernel Flow1 Time Domain functional Near-infrared Spectroscopy (TD-fNIRS)^26^. Subjective effects of alcohol were assessed with the Biphasic Alcohol Effects Scale (BAES)^27^.

Upon arrival for all study visits, participants were breathalyzed (Intoxilyzer 800, CMI, Inc, Owensboro, KY) to ensure a 0.00 BAC. For all breathalyzer measurements, breath alcohol content (BrAC) was measured and reported as equivalent BAC. Weight was measured at each visit to calculate the appropriate alcohol dose. Participants were asked to verbally confirm that they had fasted for 3 hours, had refrained from alcohol for 24 hours, were not taking any medications that were contraindicated with alcohol, and had arranged a ride home before the session started. Participants of child-bearing potential were required to take a urine pregnancy test if there was not a negative pregnancy test on record within 7 days of all study visits. At the initial visit, age was verified with a photo ID and participants completed a demographics questionnaire including handedness along with the Questionnaire Measure of Habitual Alcohol Use^43^. Also at the initial visit, a colorimeter (Cortex Technology DSM III Color Meter, CyberDerm, Broomall, PA) was used to record melanin levels (Supplementary Fig. 7). Measurements were taken on the temporal, occipital, and parietal regions of the scalp, as well as on the forehead. Only data from the forehead readings were used for two reasons: 1) to verify participant self-report measures of skin color (Fitzpatrick Scale^44^), and 2) to determine how melanin levels affected the percentage of retained channels in the prefrontal regions of the Flow1 headset (see *<u>Data preprocessing and feature extraction</u>* of Methods for further details).

Participants were trained on the Go/No-go task and practiced the task on their first study visit with 2 example blocks, one with go-only trials and one with both go and no-go trials. At each visit, participants completed a baseline run of the Go/No-go task with the Flow1 headset before receiving a study beverage. They also completed a resting state task (data not reported).

Alcohol dosing was based on Widmark’s Equation^45^ which incorporates participant weight and sex. Alcohol was administered as vodka mixed with juice (either orange or cranberry) to total 400 mL of liquid. The beverage was equally divided into two red solo cups with lids and straws. Participants had to consume the first portion within 3 minutes. This was followed by a 1 minute break. After the break, they had to consume the second portion within 3 minutes. Participants were instructed to pace their drinking to fill the 3 minutes. In total they consumed both portions within 7 minutes. Participants received roughly 1 shot (mean±s.d.=1.33±0.38, range=1.43) of alcohol at the low dose visit and 3 shots (mean±s.d.=2.64 ± 0.76, range=2.80) at the medium dose visit. A shot is defined as 44 mL of 80 proof vodka, or 1.5 oz. At the placebo visit, the instructions remained the same, however participants received 400 mL of juice entirely and the cup lid was sprayed with vodka for blinding purposes.

After finishing the drinks, participants were breathalyzed every 5 minutes starting at 10 minutes. During this time participants were allowed to read or watch a show. One of the following had to occur before starting the second set of the Go/No-go task: 1) a participant reached a BAC of 0.03 for low dose or 0.07 for moderate dose sessions, 2) BAC peaked somewhere under 0.03 or 0.07 and went down in 2 successive measurements or 3) 30 minutes passed if 1 or 2 did not occur. For the placebo dose, participants were breathalyzed twice, at 10 and 15 minutes after finishing the drink, before returning to the task. The mean time-to-starting the second brain recording segment was 21.67 minutes for the moderate dose and 18.65 minutes for the low dose.

Before beginning the second set of tasks, participants completed the BAES with an additional question asking to report any tobacco usage in the 2 hours before the study visit (none was reported for any participant at any visit). The Flow1 headset was then placed and participants completed the Go/No-go task and a resting state task (resting state data not reported). After the second Go/No-go task, participants completed the BAES once again with an additional question asking how many shots of alcohol they thought they received that day. Participants were required to wait onsite for 30 mins before they were allowed to leave with their pre-arranged transportation, which did not include driving or riding a non-motorized transportation. Participants were offered food and beverages during the wait period and were breathalyzed at the end of the visit.

### Go/No-go task paradigm

The task was coded and presented using PsychoPy. The task consisted of two block types: go-only and go-no-go. There were 10 blocks total, with 5 go-only blocks alternating with 5 go-no-go blocks, starting with a go-only block. Each block consisted of 24 trials. Go trials were cued with green leaf cartoon images, and no-go trials were cued with red flower images, with 7 different images of each type. Stimuli were presented in a pseudorandom order which was pre-set and unique for each run. The occurrence probability of a no-go stimulus in go-no-go blocks was 19% in accordance with best practices for Go/No-go tasks^46^. The background display was a pleasant image of mountains in muted colors in order to maintain interest in the task. 6 unique pre-set versions of a task run were employed in the study, so participants never repeated the same version. All participants completed the same versions in the same order, but due to the random assignment of dose order, the versions were spread across doses equally.

The task began with a 30 second baseline resting period. At the beginning of each block, an instruction screen was visible for 3 seconds and reminded the participant when to respond with a space bar press (green leaf) and when to refrain from pressing (red flower). Stimuli were presented for 400 ms, after which a space bar response would be considered a late response for a go trial or a correct rejection for a no-go trial. Negative and positive feedback was provided through the use of two sounds, either a pleasant tone for a hit or a negative tone for a false alarm, in order to encourage responses in the optimal time window^46^. Both sounds played immediately after a space bar press and lasted less than 300 ms. Sounds were found not to interfere with the neural response during piloting of the task. No auditory feedback was provided for a correct no-go trial or a missed go trial. After the 400 ms stimulus, there was a 700 ms inter-trial interval, during which the stimulus disappeared and only the mountain background was displayed. Starting after the first block, following all 24 trials, there was a rest period of 10±2s, during which a white fixation cross was displayed over the background. The instruction screen was then displayed before the next block commenced. At the end of the last block there was a final rest period of 20 seconds to limit motion effects, during which the white fixation cross was displayed. Participants were shown an overall score at the end of the task that combined reaction time, number of consecutive correct responses, and accuracy^47^. Participants were told that the score was purely for self-comparison purposes. Completion of the task required approximately 7 minutes.

### fNIRS data collection and analyses

#### 1. Data acquisition

We used Kernel Flow1 TD-fNIRS system (Flow1, Kernel, Culver City, CA, USA) ^26,29^ to collect distributions of the times of flight of photons (DTOFs) from more than 2000 channels across the head. The system uses two wavelengths (690 nm and 850 nm) and samples at an effective rate of 7.14. Throughout the headset, there are 52 modules (consisting of a source and six detectors located equidistantly 10mm away from the source). These modules are arranged to provide coverage over prefrontal, parietal, temporal, and occipital regions across both hemispheres. Data from two participants during a single study visit was lost due to technical difficulties.

#### 2. Data preprocessing and feature extraction

Data preprocessing steps are described at length in prior work^29^. We first removed channels that did not meet criteria for carrying reliable hemodynamic and physiological signals. The number of usable channels varied based on the quality of coupling between modules and participants’ heads. However, the percentage of retained prefrontal channels (the primary region considered in this paper) fell between 80-100% and is reported for the participants when considering sex, skin type, hair color, or hair texture (Supplementary Fig. 7c).

Next, DToFs underwent dimension reduction by considering their zeroth, first and second moments (i.e. sum, mean, and variance)^48,49^. DTOF moments were corrected for motion using two sequential algorithms: Temporal Derivative Distribution Repair^50^ and cubic spline interpolation^51^ (on remaining artifacts). Motion correction was followed by a detrending step (moving average with a 100-second kernel) for epoched analyses, but not for GLM analyses (as GLM already accounts for detrending with additional regressors; see below). For our supplementary analyses using moments, an additional step of short channel regression was performed, i.e. the signals from the short-source-detector (SDS) separation channels were averaged and regressed from the longer SDS channels^52^ to remove superficial physiological signals from the cortical signal of interest.

To convert the preprocessed DToF moments data to relative changes in oxy- and deoxy-hemoglobin concentrations (HbO and HbR, respectively), we used a sensitivity method described in detail previously^29,53–55^ to obtain absorption coefficients at each wavelength. Finally, these coefficients were converted to relative changes in the two chromophores using the extinction coefficients for the two wavelengths and the modified Beer–Lambert law (mBLL)^56^. HbT was computed by summing HbO and HbR.

Lastly, only channels that were formed between sources and detectors both within the prefrontal regions were considered for subsequent analyses.

#### 3. Generalized Linear Model (GLM) approach

The GLM consists in trying to explain each of the measured time courses **y** (here, relative concentrations of HbO and HbR for each channel) with a set of regressors which form the design matrix **X**; the multiple linear regression problem can be written as y = X**β** + ***ε***, where the **β** coefficients represent the contribution of each regressor to the observed data and ***ε*** are the residuals, and is solved using a least-squares solution that includes prewhitening the time courses with an autoregressive model. The design matrix **X** includes terms that captured information about the timing of the blocks (where blocks are modeled as square waves, convolved with a canonical hemodynamic response function; one regressor for go blocks and one regressor for go-no-go blocks), as well as additional nuisance regressors (a combination of drift and low frequency cosine terms). Contrasts, and their significance (p-value) can be obtained between conditions of interest via operations on the **β** coefficients (Fig. 3).

#### 4. Lateralization metrics

This data underwent further processing, namely a filtering step (cutoff frequency of 0.1Hz). We then epoched data for each session by averaging the activity over a time interval of −5 to 45 seconds (around block start time) for each condition (go and go-no-go) separately. Time courses were then averaged across all prefrontal channels independently for each hemisphere yielding right and left activation (Fig. 4).

a. Direction metric: this metric was computed by subtracting the right and left time-averaged activation for both go-only and go-no-go blocks.
b. Angle and magnitude metrics: Here, we first defined axes in a two-dimensional plane consisting of the left (x) and right (y) prefrontal activity. The trajectory of this parametric curve in time was then quantified as follows: each time point was treated as a vector described by a magnitude and an angle. The average vector was then computed and summarized by its resulting magnitude and angle yielding the final metrics. We only considered the temporal dynamics up until the peak activation of right for go-no-go blocks and left for go-only blocks, because the parametric curves intersected with themselves beyond the peaks. To account for artifacts and atypical activation that may result in early peaks we considered at a minimum the full block duration (30 sec). These parametric representation of right and left coactivation in the described two-dimensional plane encapsulate complementary information about the covariation between right and left prefrontal activity, namely the angle metric captures a granular time-locked measure of lateralization while the magnitude metric contains information about the absolute change in prefrontal activation.

#### 5. Placebo-effect analyses

Placebo-effect was inferred from participants’ self-reported estimate of the number of standard shots contained in their drink that session. For placebo dosing sessions, if participants indicated that they received any amount of alcohol, we consider them to have had a placebo-effect, and those reporting zero standard shots we considered to have had no placebo-effect. Two participants were excluded from this analysis due to missing subjective reports at the placebo study visit. To compare the angle metric between those with (n=23) and without (n=23) placebo-effect, we used the Watson Williams test from Python’s pycircstat package^57^.

#### 6. Linear Models

We considered a series of Ordinary Least Squares (OLS) regression models to examine the relationship between post-dose behavior (d-prime in go-no-go blocks) as a function of different input variables (Fig. 5). To test the performance of the models, two metrics were used: (1) R-squared values: indicative of model’s goodness of fit. (2) Akaike information criterion (AIC): AIC = 2k-2ln(L), where k and L indicate the number of parameters and maximum log likelihood respectively. This measure is indicative of model improvement while accounting for overfitting after introducing additional variables. Moreover, it can only be interpreted as a comparison metric, and not a stand alone metric.

We compared the performance of the four following models:

1. Basic model: Input variables included pre-dose behavior, sex (0:female/1:male) and age of the participant (3 total variables).
2. BAC model: Direct measure of intoxication (BAC) was included as an input variable in addition to those in the basic model (4 total variables).
3. Direction model: The direction lateralization metric was added as a variable to those in the BAC model (5 total variables)
4. Angle model: To account for the circular nature of the angle metric, and to include this measure as a variable in our linear models, we computed the sine and cosine of the angles, multiplied by the corresponding magnitudes separately. These two variables were simultaneously included as input variables in addition to those in the BAC model (6 total variables).

To control for family-wise error rate, we used a Bonferroni correction to obtain adjusted p-values when comparing the four models.

A separate additional model was used to investigate the role of subjective measures of intoxication.

## Acknowledgments

We thank Adelaida Castillo for regulatory support for this study.

## Author contributions

Conceptualization: KLP, RMF. Methodology/Software: JD, ZMA, EMK, SJ. Formal analysis: EMK, ZMA, MT, NM. Investigation: MT, NM, AJ, EMK. Writing - Original Draft. EMK, ZMA, MT, NM, KLP. Writing - Review and Editing: all authors. Author order was determined alphabetically. Dr. Katherine Perdue agrees to be accountable for all aspects of the work, ensuring that questions related to the accuracy or integrity of any part of the work are appropriately investigated and resolved.

## Data availability statement

The datasets used and/or analyzed during the current study are available from the corresponding author on reasonable request.

## Supplementary Material

**Supplementary Figure 1:**
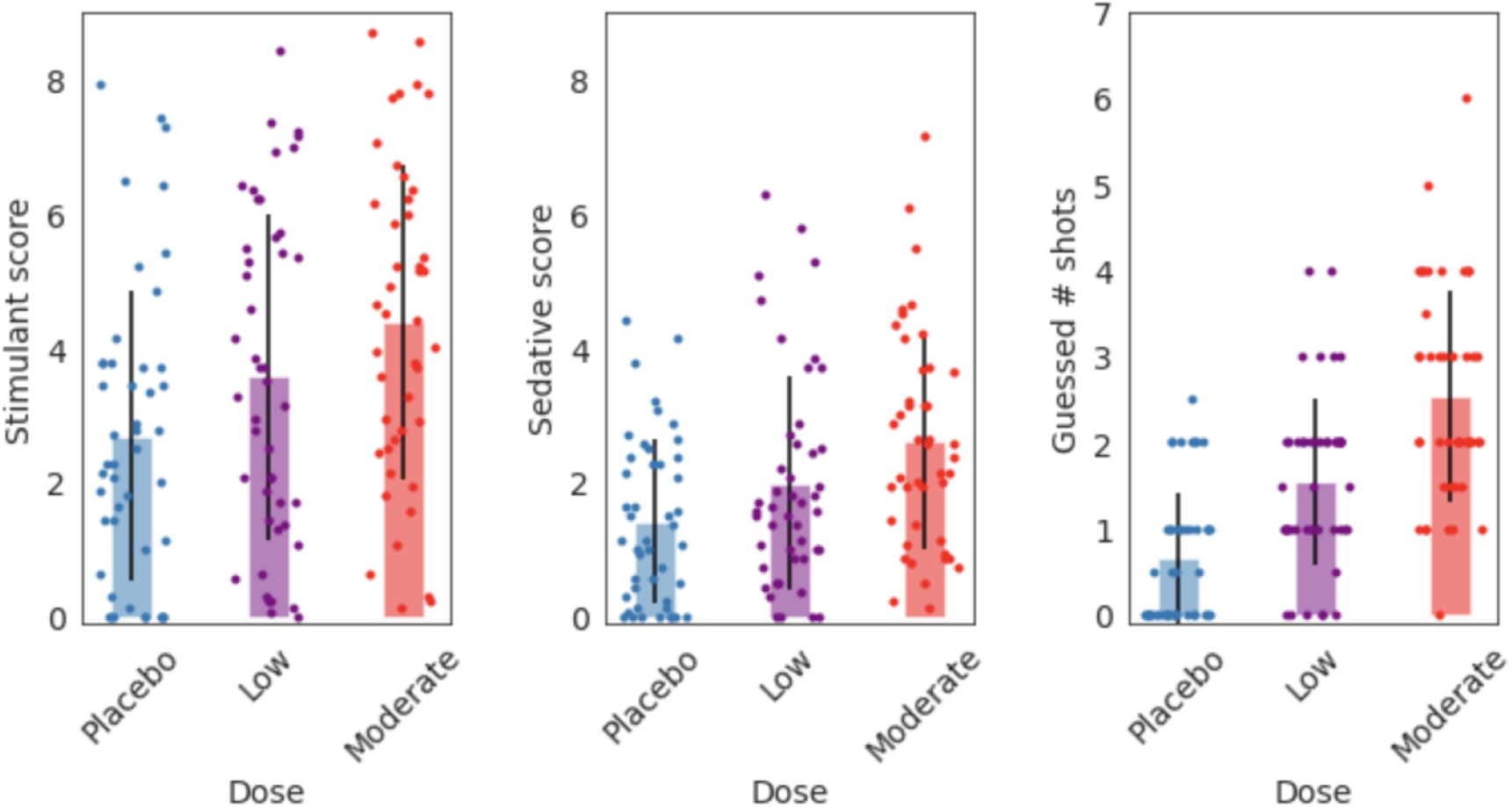
Subjective measures of intoxication. Shown are the subjective measures of intoxication for each participant (each dot) at different doses. For each measure, there was a significant difference between all dose pairs. Stimulant scores (left) - paired t-test p-values: placebo/low = 9.21×10^−3^, placebo/moderate = 1.49×10^−5^, low/moderate = 1.50×10^−3^); Sedative score (middle) - paired t-test p-values: placebo/low = 0.019, placebo/moderate = 1.32×10^−5^, low/moderate = 9.98×10^−3^); Guessed number of shots (right): paired t-test p-values: placebo/low = 3.31×10^−5^, placebo/moderate = 2.60×10^−11^, low/moderate = 5.18×10^−7^)

**Supplementary Figure 2:**
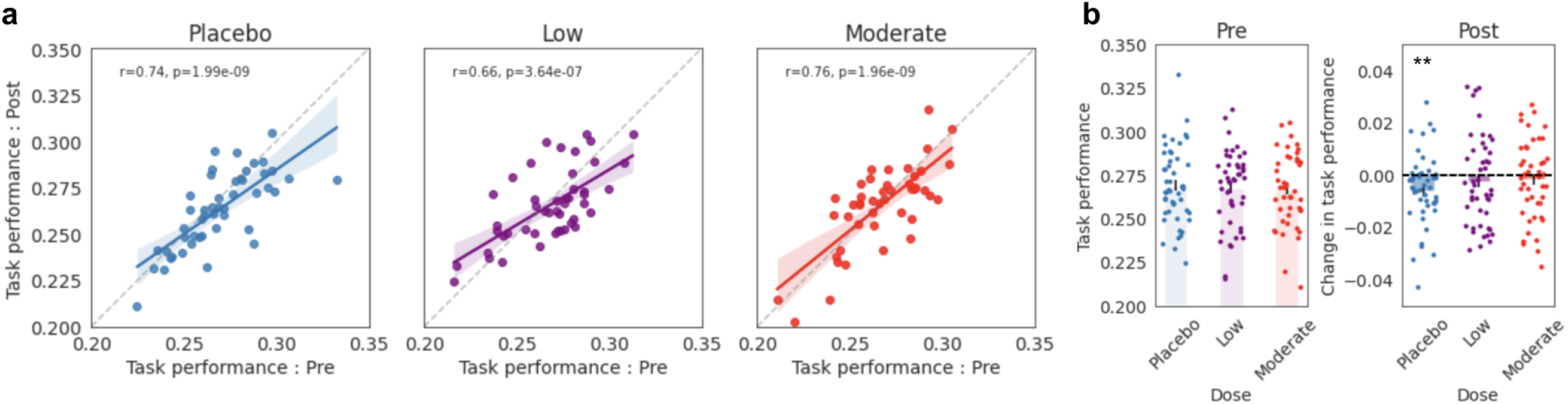
Main effect of alcohol level on task performance as measured by reaction times. **a.** Pre Go/No-go performance (reaction time measured in seconds) is plotted against post Go/No-go performance (reaction time measured in seconds) for placebo (left, blue), low (middle, purple), and moderate (right, red) levels of intoxication. Solid lines depict the line-of-best-fit and the shaded regions show the 95% confidence intervals. Dashed lines indicate the line of unity. Task performance is computed as the reaction times in the go-no-go blocks. **b.** Left: Colored bars show the average reaction times across the pre tasks for placebo, low, and moderate sessions separately. Right: The change in performance, measured as post reaction time - pre reaction time are shown for the different doses. Colored dots show individual subject measures (mean±s.e.m). Reaction times were significantly reduced in the post task compared to the pre task only for the placebo dose (paired t-test p=9.45×10^−3^).

**Supplementary Figure 3:**
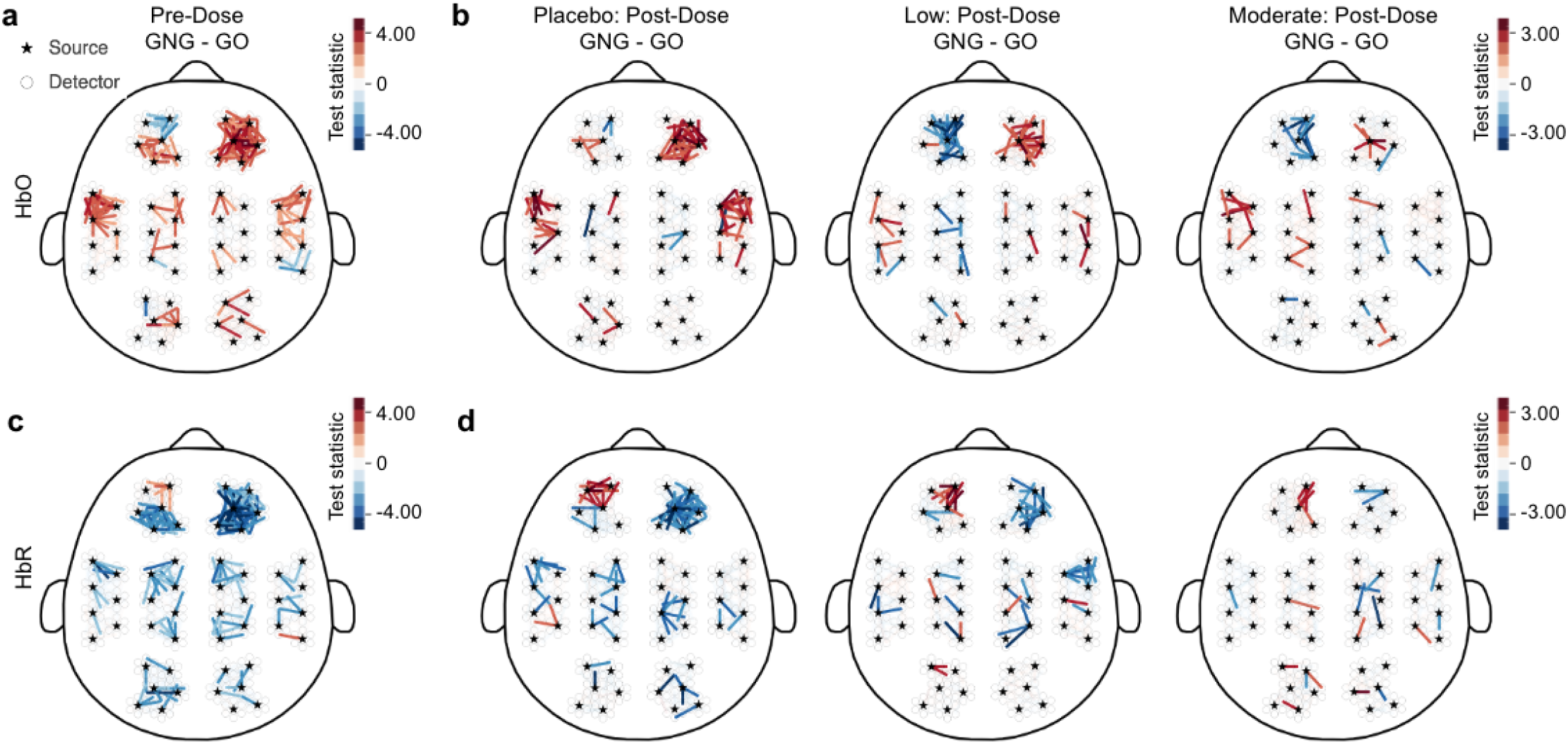
Whole-head brain activation during an inhibitory control task under the influence of different alcohol doses as quantified by generalized linear models. **a-b.** The contrast between brain activity (measured via HbO chromophore) during go-no-go blocks compared to the go-only blocks in pre-dose, and post-placebo, post-low, and post-moderate sessions (increased activation is indicated by warmer colors and decreased activation is indicated by cooler colors). Each line is an individual channel and only significant channels at p<0.05 are displayed. **c-d.** Same as (a-b) but for HbR. In all subpanels, the color scale indicates the t-statistic.

**Supplementary Figure 4:**
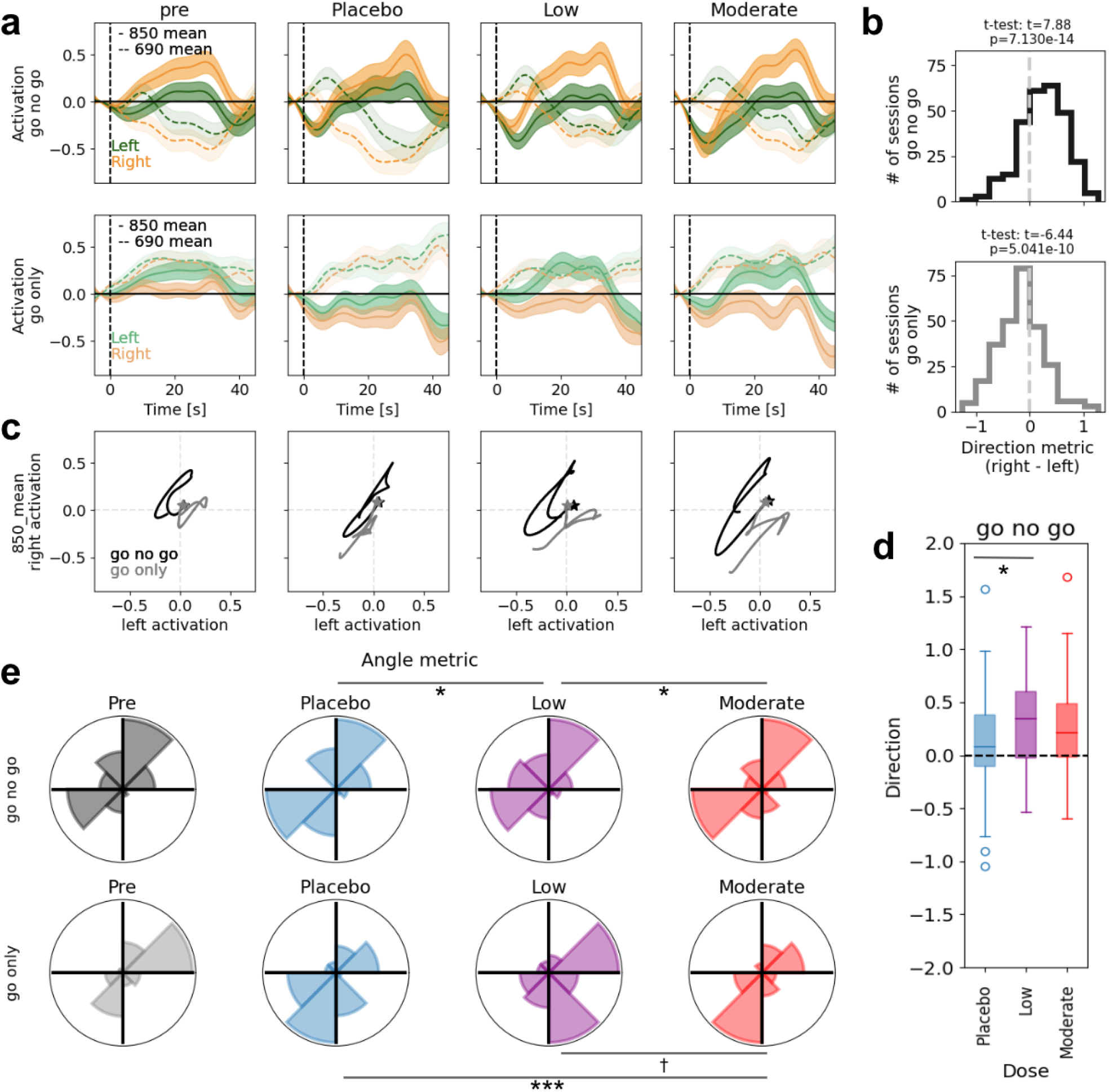
DToF mean moment of 850nm shows altered patterns of prefrontal lateralization with different alcohol doses. **a.** Block averaged time course of left (green) and right (orange) prefrontal activity (-mean moments) for go-no-go blocks (top) and go-only blocks (bottom) separated by session. From left to right, pre-dose and placebo-, low-, and moderate-dose time courses are shown (mean±s.e.m). Solid and dashed lines represent the 850nm and 690nm wavelengths mean moment respectively. Note the opposite trends in the two wavelengths, as expected. **b.** Population distribution of lateralization (direction metric) for go-no-go (top; black) and go-only (bottom; gray) blocks across all participants and sessions. Note the right lateralization for go-no-go blocks (shift towards positive values, t-test: t = 7.88, p=7.13×10^−14^) and left lateralization for go-only blocks (shift towards negative values, t-test: t=-6.44, p=5.04×10^−10^). **c.** Population average parametric curves showing the trajectory of right prefrontal vs. left prefrontal activity through time for go-only (gray) and go-no-go (black) blocks, split by pre-dose and session type, as in (a). Stars indicate initial time points. **d.** The distributions of the direction metric for go-no-go blocks across post-dose sessions (* significant difference between placebo and low dose: p<0.05). **e.** The distributions of the angle metric, split by pre-dose and session type, as in (a). * and *** indicate significant deviation from uniformity (at *p* < 0.05 and p < 0.001 respectively; rayleigh test; † trend towards significant) when computing pairwise difference in the angle metric between dosing sessions.

**Supplementary Figure 5:**
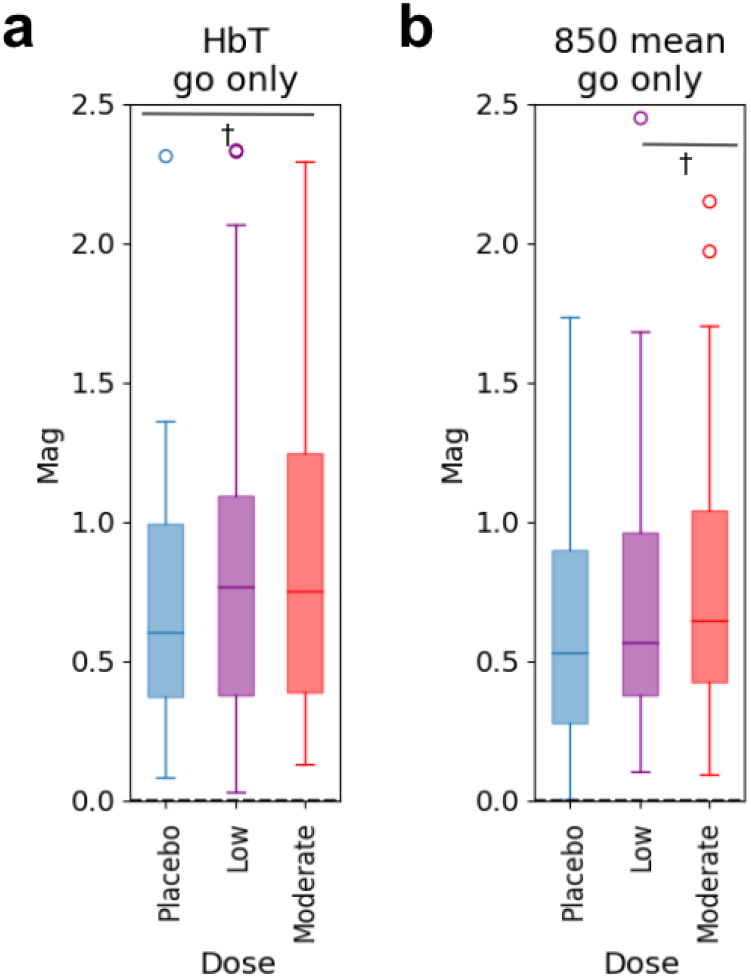
Dose-dependent trends in magnitude metric during go-only condition of the Go/No-go Task. **a.** The distributions of the HbT magnitude metric for go-only blocks across post-dose sessions (paired t-test: † trend towards significant difference between placebo and moderate dose: p=0.06). **b.** The distributions of the DToF mean moment (of 850nm) magnitude metric for go-only blocks across post-dose sessions (paired t-test: † trend towards significant difference between low and moderate dose: p=0.07).

**Supplementary Figure 6:**
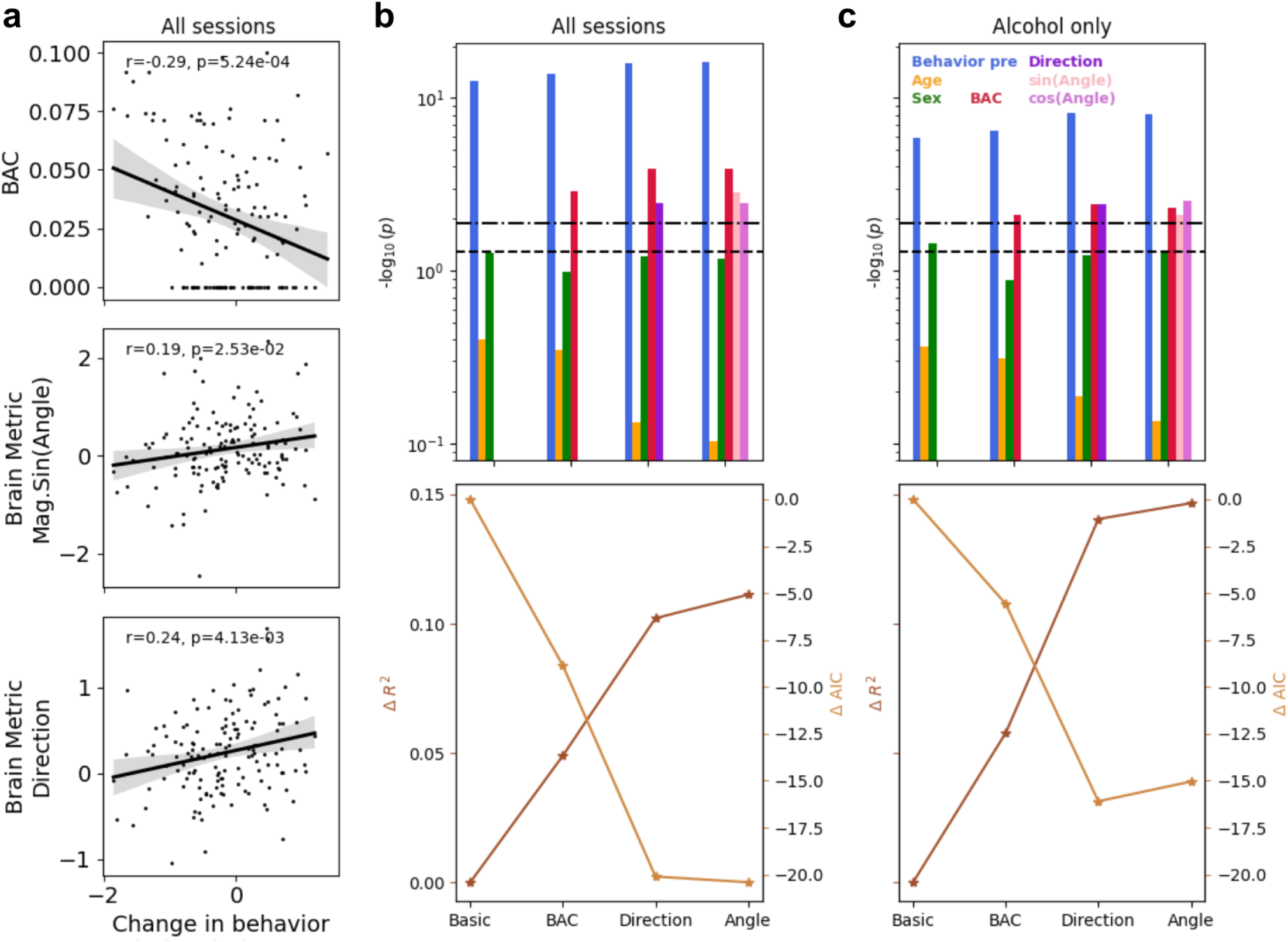
DToF mean moment of 850nm lateralization metrics improved the prediction of behavioral performance in addition to measured BAC. **a.** The change in behavioral performance as measured by post d-prime - pre d-prime was significantly correlated with BAC (top; pearson r=-0.29, p=5.24×10^−4^), magnitude x sin(angle) (middle; pearson r=0.19, p=2.53×10^−2^), and direction metric (bottom; pearson r=0.24, p=4.13×10^−3^). **b.** Top) Statistical significance of the different factors in the four OLS models (four blocks shown on the x-axis) in predicting the post behavioral performance are reported as −log_10_(p-value). We build off of the three factors in the basic model (pre behavioral performance: blue, age: yellow, sex: green), and include additional variables in separate OLS models: 1) BAC model included the direct measure of intoxication (BAC values, red) in addition to the basic model; 2) Direction model consisted of the direction metric (dark purple) in addition to the BAC model variables; 3) Angle model included the weighted sine and cosine of the angle metric (pink and light purple respectively) in addition to the BAC model variables. Note that both BAC values and lateralization metrics were significant factors in predicting post performance. Dashed lines indicate a significance level of 0.05 and 0.05/4 (after correcting for multiple comparisons using Bonferroni correction; Methods). Bottom) Overall OLS model performances are shown in terms of the change in R-squared values and the change in AIC (lower AIC values indicate better performance). **c.** Same as (b) but when only considering the sessions with alcohol consumption (i.e. low and moderate doses).

**Supplementary Figure 7:**
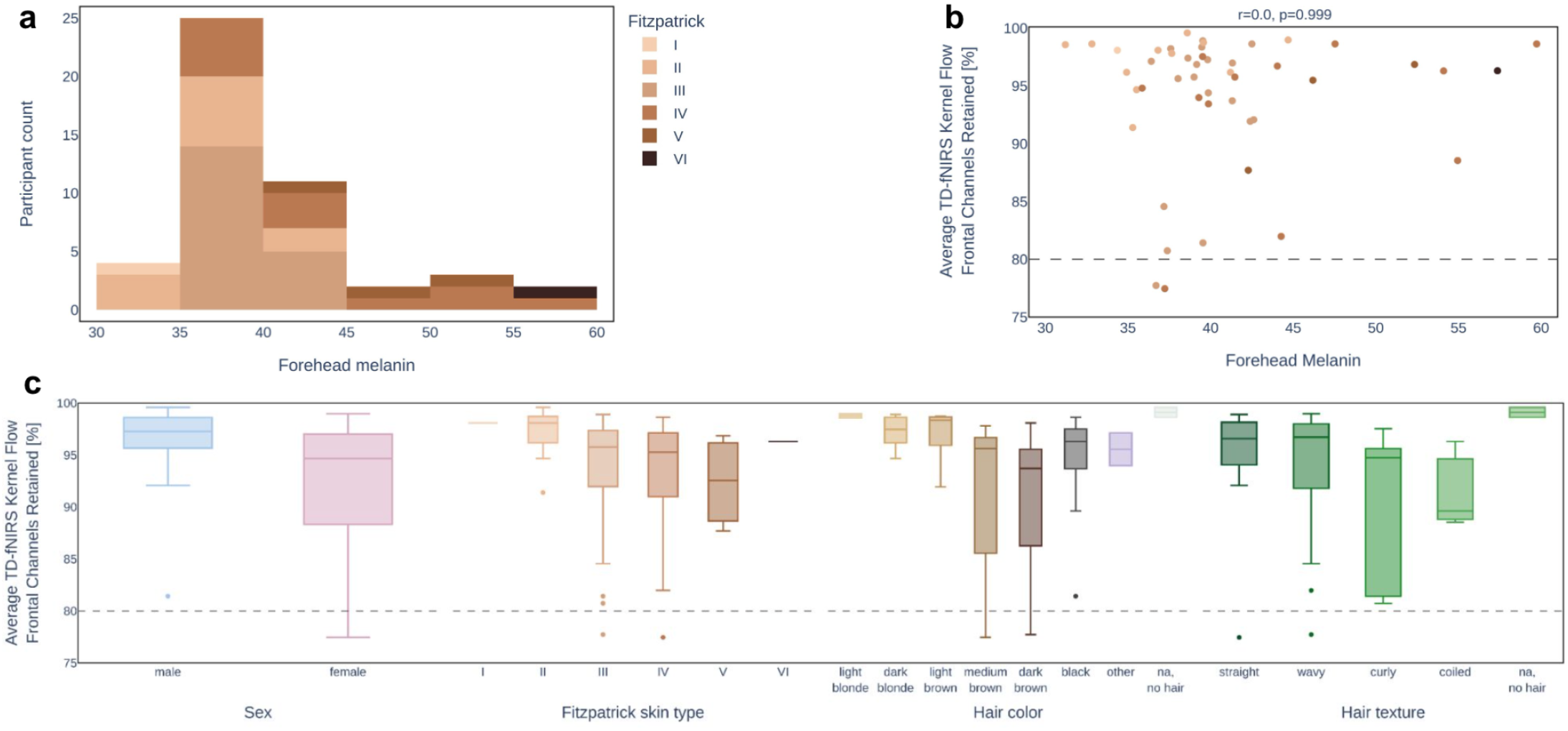
The relationship between participants’ skin color and the percentage of retained channels. **a.** A histogram of participants’ forehead melanin levels, as measured by the colorimeter. Histogram is shaded by self-reported Fitzpatrick skin type (scale of I to VI, I being fairer and VI being darker skin). The participant pool contained a range of melanin levels. **b.** The percentage of retained prefrontal channels as a function of recorded forehead melanin levels. Each point represents a participant with the shade representing the participant’s self-reported Fitzpatrick skin type (see a.). All but 2 participants had greater than 80% of channels retained (seen as the points above the dashed line), and all were above 75%. A lack of correlation indicated that the Flow1 system was able to perform well on a variety of skin colors. **c.** The distributions of retained prefrontal channels separated by various demographic factors (sex, Fitzpatrick skin type, hair color, and hair texture).

**Supplementary Table 1:** Participant Demographics and smoking history.

|  | Male (n=23) | Female (n=25) | All (n=48) |
| --- | --- | --- | --- |
| Age | 32.48 ± 9.08 | 32.76 ± 10.59 | 32.63 ± 9.90 |
| Mean drinks per week | 6.24 ± 2.79 | 4.76 ± 2.45 | 5.43 ± 2.72 |

**Supplementary Table 1: Participant Demographics and smoking history.**
|  | All (n=48) |
| --- | --- |
| Handedness | right: n=32<br>mixed: n=11<br>left: n=5 |
| Ever smoked | n=15 |
| Mean years smoked | 4 (n=13, 2 preferred not to answer) |
| Mean cigarettes smoked/day | 3.92 (n=12, 3 preferred not to answer) |

## References

1. Zoethout, R. W. M., Delgado, W. L., Ippel, A. E., Dahan, A. & van Gerven, J. M. A. Functional biomarkers for the acute effects of alcohol on the central nervous system in healthy volunteers: Functional biomarkers for the acute CNS effects of alcohol. British Journal of Clinical Pharmacology 71, 331–350 (2011).

2. Field, M., Wiers, R. W., Christiansen, P., Fillmore, M. T. & Verster, J. C. Acute Alcohol Effects on Inhibitory Control and Implicit Cognition: Implications for Loss of Control Over Drinking. Alcoholism: Clinical and Experimental Research no-no (2010) doi:10.1111/j.1530-0277.2010.01218.x.

3. Moskowitz, H. & Fiorentino, D. A review of the literature on the effects of low doses of alcohol on driving-related skills: (441302008-001). (2000) doi:10.1037/e441302008-001.

4. Fillmore, M. T. Acute alcohol-induced impairment of cognitive functions: Past and present findings. International Journal on Disability and Human Development 6, (2007).

5. Abroms, B. D., Fillmore, M. T. & Marczinski, C. A. Alcohol-induced impairment of behavioral control: effects on the alteration and suppression of prepotent responses. J. Stud. Alcohol 64, 687–695 (2003).

6. Fillmore, M. T., Blackburn, J. S. & Harrison, E. L. R. Acute disinhibiting effects of alcohol as a factor in risky driving behavior. Drug and Alcohol Dependence 95, 97–106 (2008).

7. George, W. H. & Stoner, S. A. Understanding acute alcohol effects on sexual behavior. Annu Rev Sex Res 11, 92–124 (2000).

8. Ito, T. A., Miller, N. & Pollock, V. E. Alcohol and Aggression: A Meta-Analysis on the Moderating Effects of Inhibitory Cues, Triggering Events, and Self-Focused Attention. Psychological Bulletin 120, 60–82 (1996).

9. Donders, F. C. On the speed of mental processes. Acta Psychologica 30, 412–431 (1969).

10. Fillmore, M. T., Marczinski, C. A. & Bowman, A. M. Acute tolerance to alcohol effects on inhibitory and activational mechanisms of behavioral control. J. Stud. Alcohol 66, 663–672 (2005).

11. Gan, G. et al. Alcohol-Induced Impairment of Inhibitory Control Is Linked to Attenuated Brain Responses in Right Fronto-Temporal Cortex. Biological Psychiatry 76, 698–707 (2014).

12. Rubia, K. et al. Mapping Motor Inhibition: Conjunctive Brain Activations across Different Versions of Go/No-Go and Stop Tasks. NeuroImage 13, 250–261 (2001).

13. Garavan, H., Ross, T. J. & Stein, E. A. Right hemispheric dominance of inhibitory control: An event-related functional MRI study. Proc. Natl. Acad. Sci. U.S.A. 96, 8301–8306 (1999).

14. Simmonds, D. J., Pekar, J. J. & Mostofsky, S. H. Meta-analysis of Go/No-go tasks demonstrating that fMRI activation associated with response inhibition is task-dependent. Neuropsychologia 46, 224–232 (2008).

15. Bjork, J. M. & Gilman, J. M. The effects of acute alcohol administration on the human brain: Insights from neuroimaging. Neuropharmacology 84, 101–110 (2014).

16. Claus, E. D. & Hendershot, C. S. Moderating effect of working memory capacity on acute alcohol effects on BOLD response during inhibition and error monitoring in male heavy drinkers. Psychopharmacology 232, 765–776 (2015).

17. Gundersen, H., Specht, K., Grüner, R., Ersland, L. & Hugdahl, K. Separating the effects of alcohol and expectancy on brain activation: An fMRI working memory study. NeuroImage 42, 1587–1596 (2008).

18. Anderson, B. M. et al. Functional Imaging of Cognitive Control During Acute Alcohol Intoxication. Alcoholism: Clinical and Experimental Research 35, 156–165 (2011).

19. Nikolaou, K., Critchley, H. & Duka, T. Alcohol Affects Neuronal Substrates of Response Inhibition but Not of Perceptual Processing of Stimuli Signalling a Stop Response. PLoS ONE 8, e76649 (2013).

20. Tsujii, T., Sakatani, K., Nakashima, E., Igarashi, T. & Katayama, Y. Characterization of the acute effects of alcohol on asymmetry of inferior frontal cortex activity during a Go/No-Go task using functional near-infrared spectroscopy. Psychopharmacology 217, 595–603 (2011).

21. Carollo, A., Cataldo, I., Fong, S., Corazza, O. & Esposito, G. Unfolding the real-time neural mechanisms in addiction: Functional near-infrared spectroscopy (fNIRS) as a resourceful tool for research and clinical practice. Addiction Neuroscience 4, 100048 (2022).

22. Heilbronner, U. & Münte, T. F. Rapid event-related near-infrared spectroscopy detects age-related qualitative changes in the neural correlates of response inhibition. NeuroImage 65, 408–415 (2013).

23. Cui, X., Bray, S., Bryant, D. M., Glover, G. H. & Reiss, A. L. A quantitative comparison of NIRS and fMRI across multiple cognitive tasks. NeuroImage 54, 2808–2821 (2011).

24. Herrmann, M. J., Plichta, M. M., Ehlis, A.-C. & Fallgatter, A. J. Optical topography during a Go–NoGo task assessed with multi-channel near-infrared spectroscopy. Behavioural Brain Research 160, 135–140 (2005).

25. Obata, A., Morimoto, K., Sato, H., Maki, A. & Koizumi, H. Acute effects of alcohol on hemodynamic changes during visual stimulation assessed using 24-channel near-infrared spectroscopy. Psychiatry Research: Neuroimaging 123, 145–152 (2003).

26. Ban, H. Y. et al. Kernel Flow: a high channel count scalable time-domain functional near-infrared spectroscopy system. J. Biomed. Opt. 27, (2022).

27. Martin, C. S., Earleywine, M., Musty, R. E., Perrine, M. W. & Swift, R. M. Development and Validation of the Biphasic Alcohol Effects Scale. Alcoholism Clin Exp Res 17, 140–146 (1993).

28. Miller, M. A. & Fillmore, M. T. Protracted impairment of impulse control under an acute dose of alcohol: A time-course analysis. Addictive Behaviors 39, 1589–1596 (2014).

29. Castillo, A. et al. Acute effects of subanesthetic ketamine on cerebrovascular hemodynamics in humans: A TD-fNIRS neuroimaging study. Preprint at https://doi.org/10.1101/2023.01.06.522912.

30. Gagnon, L. et al. Quantification of the cortical contribution to the NIRS signal over the motor cortex using concurrent NIRS-fMRI measurements. NeuroImage 59, 3933–3940 (2012).

31. Abdalmalak, A. et al. Can time-resolved NIRS provide the sensitivity to detect brain activity during motor imagery consistently? Biomed. Opt. Express 8, 2162 (2017).

32. Torricelli, A. et al. Time domain functional NIRS imaging for human brain mapping. NeuroImage 85, 28–50 (2014).

33. Milej, D. et al. Time-resolved multi-channel optical system for assessment of brain oxygenation and perfusion by monitoring of diffuse reflectance and fluorescence. Opto-Electronics Review 22, (2014).

34. Monden, Y. et al. Right prefrontal activation as a neuro-functional biomarker for monitoring acute effects of methylphenidate in ADHD children: An fNIRS study. NeuroImage: Clinical 1, 131–140 (2012).

35. Konishi, S. et al. Common inhibitory mechanism in human inferior prefrontal cortex revealed by event-related functional MRI. Brain 122, 981–991 (1999).

36. Wager, T. D. et al. Common and unique components of response inhibition revealed by fMRI. NeuroImage 27, 323–340 (2005).

37. Schaefer, A. et al. Local-Global Parcellation of the Human Cerebral Cortex from Intrinsic Functional Connectivity MRI. Cerebral Cortex 28, 3095–3114 (2018).

38. Weafer, J., Phan, K. L. & de Wit, H. Poor inhibitory control is associated with greater stimulation and less sedation following alcohol. Psychopharmacology 237, 825–832 (2020).

39. Weafer, J. et al. Neural correlates of inhibitory control are associated with stimulant-like effects of alcohol. Neuropsychopharmacol. 46, 1442–1450 (2021).

40. Johnstone, L. T., Karlsson, E. M. & Carey, D. P. Left-Handers Are Less Lateralized Than Right-Handers for Both Left and Right Hemispheric Functions. Cerebral Cortex bhab048 (2021) doi:10.1093/cercor/bhab048.

41. Ewing, J. A. Detecting Alcoholism: The CAGE Questionnaire. JAMA 252, 1905–1907 (1984).

42. Saunders, J. B., Aasland, O. G., Babor, T. F., De La Fuente, J. R. & Grant, M. Development of the Alcohol Use Disorders Identification Test (AUDIT): WHO Collaborative Project on Early Detection of Persons with Harmful Alcohol Consumption-II. Addiction 88, 791–804 (1993).

43. Mehrabian, A. & Russell, J. A. A Questionnaire Measure of Habitual Alcohol Use. Psychol Rep 43, 803–806 (1978).

44. Fitzpatrick, T. B. The Validity and Practicality of Sun-Reactive Skin Types I Through VI. Arch Dermatol 124, 869 (1988).

45. Roberts, C. & Robinson, S. P. Alcohol concentration and carbonation of drinks: The effect on blood alcohol levels. Journal of Forensic and Legal Medicine 14, 398–405 (2007).

46. Wessel, J. R. Prepotent motor activity and inhibitory control demands in different variants of the go/no-go paradigm. Psychophysiol 55, e12871 (2018).

47. Lumsden, J., Skinner, A., Woods, A. T., Lawrence, N. S. & Munafò, M. The effects of gamelike features and test location on cognitive test performance and participant enjoyment. PeerJ 4, e2184 (2016).

48. Wabnitz, H., Contini, D., Spinelli, L., Torricelli, A. & Liebert, A. Depth-selective data analysis for time-domain fNIRS: moments vs time windows. Biomed. Opt. Express 11, 4224 (2020).

49. Liebert, A. et al. Evaluation of optical properties of highly scattering media by moments of distributions of times of flight of photons. Appl. Opt. 42, 5785 (2003).

50. Fishburn, F. A., Ludlum, R. S., Vaidya, C. J. & Medvedev, A. V. Temporal Derivative Distribution Repair (TDDR): A motion correction method for fNIRS. NeuroImage 184, 171–179 (2019).

51. Scholkmann, F., Spichtig, S., Muehlemann, T. & Wolf, M. How to detect and reduce movement artifacts in near-infrared imaging using moving standard deviation and spline interpolation. Physiol. Meas. 31, 649–662 (2010).

52. Gagnon, L. et al. Improved recovery of the hemodynamic response in diffuse optical imaging using short optode separations and state-space modeling. NeuroImage 56, 1362–1371 (2011).

53. Ortega-Martinez, A. et al. How much do time-domain functional near-infrared spectroscopy (fNIRS) moments improve estimation of brain activity over traditional fNIRS? Neurophoton. 10, (2022).

54. Dehghani, H. et al. Near infrared optical tomography using NIRFAST: Algorithm for numerical model and image reconstruction. Commun. Numer. Meth. Engng. 25, 711–732 (2009).

55. Doulgerakis, M., Eggebrecht, A., Wojtkiewicz, S., Culver, J. & Dehghani, H. Toward real-time diffuse optical tomography: accelerating light propagation modeling employing parallel computing on GPU and CPU. J. Biomed. Opt 22, 1 (2017).

56. Huppert, T. J., Diamond, S. G., Franceschini, M. A. & Boas, D. A. HomER: a review of time-series analysis methods for near-infrared spectroscopy of the brain. Appl. Opt. 48, D280 (2009).

57. Berens, P. pycircstat Documentation. (2018).

